# Genomic adaptation of the picoeukaryote *Pelagomonas calceolata* to iron-poor oceans revealed by a chromosome-scale genome sequence

**DOI:** 10.1101/2021.10.25.465678

**Authors:** Nina Guérin, Marta Ciccarella, Elisa Flamant, Paul Frémont, Sophie Mangenot, Benjamin Istace, Benjamin Noel, Sarah Romac, Charles Bachy, Martin Gachenot, Eric Pelletier, Adriana Alberti, Olivier Jaillon, Corinne Cruaud, Patrick Wincker, Jean-Marc Aury, Quentin Carradec

## Abstract

The smallest phytoplankton species are key actors in oceans biogeochemical cycling and their abundance and distribution are affected with global environmental changes. Picoalgae (cells <2µm) of the Pelagophyceae class encompass coastal species causative of harmful algal blooms while others are cosmopolitan and abundant in open ocean ecosystems. Despite the ecological importance of Pelagophytes, only a few genomic references exist limiting our capacity to identify them and study their adaptation mechanisms in a changing environment. Here, we report the complete chromosome-scale assembled genome sequence of *Pelagomonas calceolata*. We identified unusual large low-GC and gene-rich regions potentially representing centromeres. These particular genomic structures could be explained by the absence of genes from a recombination pathway involving double Holiday Junctions. We identified a large repertoire of genes involved in inorganic nitrogen sensing and uptake and several genes replacing iron-requiring proteins potentially explaining *P. calceolata* ecological success in oligotrophic waters. Finally, based on this high-quality assembly, we evaluated *P. calceolata* relative abundance in all oceans using environmental *Tara* Oceans datasets. Our results suggest that *P. calceolata* is one of the most abundant eukaryotic species in the oceans with a relative abundance favoured by high temperature and iron-poor conditions. Climate change projections based on its relative abundance suggest an extension of the *P. calceolata* habitat toward the poles at the end of this century. Collectively, these findings reveal the ecological importance of *P. calceolata* and lay the foundation for a global scale analysis of the adaptation and acclimation strategies of picoalgae in a changing environment.

## Introduction

Marine phytoplankton account for more than 45% of photosynthetic primary production on Earth and play an essential role in supplying organic matter to marine food webs^1^. They are key global actors in CO_2_ uptake and provide gaseous oxygen to the atmosphere. A global decline of phytoplankton biomass has been reported over the past century (1% of chlorophyll-a concentration per year) leading to a decrease of net primary production in many oceanic regions^2^. This decline is probably a consequence of global ocean warming which drives water column stratification, reducing the nutrient supply to surface waters. Temperature-driven reductions in phytoplankton productivity in the tropics and temperate regions are likely to have cascading effects on higher trophic levels and ecosystem functioning^3^.

Picophytoplankton are photosynthetic unicellular organisms <2 μm in cell diameter. They encompass the abundant cyanobacteria *Synechococcus* and *Prochlorococcus* and photosynthetic picoeukaryotes (PPEs) belonging to different phyla including Chlorophyta, Cryptophyta, Haptophyta and Stramenopiles^4^. PPEs, present in all oceans, are the dominant primary producers in warm and oligotrophic regions. Ocean warming and expansion of oligotrophic regions in the next decades may extend the ecological niche of PPEs and a global shift from large photosynthetic organisms toward smaller primary producers is expected^3,5^. For example, sea ice melting in the Canadian Arctic Basin has been associated with an increase in the abundance of PPEs such as *Micromonas* at the expense of larger algae^6^. In the laboratory, this alga has the capacity to change its optimum temperature for growth in only a few hundred generations, which suggests that it will be less affected by global warming than many larger organisms^7^. In addition, the larger cell surface-to-volume ratio of PPEs compared to larger phytoplankton cells is advantageous for resource acquisition and growth in nutrient-limited environments^8,9^.

Iron is one key compound required for the activity of the respiratory chain, photosynthesis and nitrogen fixation^9^. Because bioavailable iron is extremely low in more than one third of the surface ocean, small phytoplankton have developed several strategies to optimize iron uptake and reduce iron needs^10^. In diatoms, reductive and non-reductive iron uptake mechanisms involve many proteins including phytotransferrins, transmembrane ferric reductases, iron permeases and siderophore-binding proteins^11^. The iron needs can be modulated by variation of gene expression levels between iron-required proteins and their iron-free equivalent. These protein switches include genes involved in electron transfer (flavodoxin/ferredoxin and plastocyanin/cytochrome c6), in gluconeogenesis (fructose-bisphosphate aldolase type I or type II) and superoxide dismutases (Mn/Fe-SOD, Cu/Zn-SOD or Ni-SOD)^12–14^.

PPE growth is also limited by nitrogen (N) availability in large portions of the global ocean^15^. Ammonium (NH_4_^+^), nitrate (NO_3_^−^) and nitrite (NO_2_^−^) are the main source of inorganic N for PPEs, however several studies have shown that dissolved organic N, like urea, can be metabolised in N-limited environments^16^. For example, several membrane-localized urea transporters in the diatom *Phaeodactylum tricornutum* are maximally expressed in nitrogen-limited conditions^17^ and the harmful algal blooms of the pelagophyceae *Aureococcus anophagefferens* may be fuelled by urea^18^.

Despite their large taxonomic distribution, most molecular studies on the ecological role of PPEs and their adaptation to the environment are restricted to a few species. PPEs are suspected to possess highly developed acclimation/adaptation capacities, but the underlying molecular mechanisms remain poorly characterized due to the lack of reference genomic data.

Among PPEs, *Pelagomonas calceolata* was the first described member of the heterokont class Pelagophyceae^19^. It has since been identified in many oceanic regions using its 18S rRNA sequence and chloroplast genome^20–22^. Several studies have demonstrated the capacity of *P. calceolata* to adapt to different environmental conditions. In the laboratory, *P. calceolata* has been shown to exhibit a high degree of acclimation to light fluctuations with rapid activation of the photo-protective xanthophyll cycle and non-photochemical quenching^23^. In the Marquesas archipelago, *P. calceolata* is one of the most responsive species to iron fertilization with upregulation of genes involved in photosynthesis, amino acid synthesis and nitrogen assimilation^12^. A global scale analysis of pelagophyte genes revealed that they are adapted to low iron conditions^13^. In the subtropical Pacific, *P. calceolata* expresses stress genes in surface samples and genes involved in nitrogen assimilation are overexpressed in the deep chlorophyll maximum^24^. A laboratory study suggests that *P. calceolata* also has the ability to increase the transcription levels of organic-nitrogenous compound cleavage enzymes (cathepsin, urease, arginase) under low nitrogen concentration^25^. Thus, gene expression appears to be controlled according to the nitrogen source and quantity. Taken together, this apparent adaptive plasticity may explain the presence of *P. calceolata* in many different oceanic environments, however, an exhaustive analysis of the genetic capacity of this species and the *in situ* characterization of its ecological niche is lacking.

Here we sequenced, assembled and annotated the *Pelagomonas calceolata* nuclear genome, with a combination of long- and short-reads. We examined genomic structure and gene content relative to other unicellular phytoplankton. Specific analyses were performed to gain new insights into its life cycle and its genetic capacity for nutrient uptake. Finally, we used this genome to detect *P. calceolata* in environmental datasets of *Tara* expeditions across all oceans, to characterize its ecological niche and to identify the environmental conditions controlling its relative abundance.

## Material and Methods

### Pelagomonas culture

*Pelagomonas calceolata* RCC100 culture was grown in 12:12-h light:dark photoperiod in K medium with natural seawater base at 20°C. At the Roscoff Culture Collection, cells were kept at a light intensity of ∼80 μmol photon m-2 s-1 and volume of culture was ramped up to 1 litre in mid-exponential growth phase before harvesting. RCC100 culture was not axenic and grown in the presence of undefined bacterial microbiota.

### DNA extraction, library preparation and sequencing

We pelleted cells from 500 ml of culture by two successive centrifugations at 10,000 g for 15 minutes at 4°C. Genomic DNA was extracted using the NucleoSpin Plant II Mini kit according to the manufacturer’s instructions (Macherey-Nagel, Hoerdt, France) with the following exception for the lysis step : 400µL of lysis buffer PL1 and 25µL of proteinase K 25mg/mL were added to strain pellets, and lysates were incubated at 55°C for 1 hour at 900 rpm. DNA quantity and integrity were respectively evaluated on a Qubit 2.0 spectrofluorometer (Invitrogen, Carlsbad, CA, USA) and a Nanodrop1000 spectrophotometer (Thermo Fisher Scientific, MA, USA).

For Illumina sequencing, DNA (1.5μg) was sonicated using a Covaris E220 sonicator (Covaris, Woburn, MA, USA). Fragments were end-repaired, 3′-adenylated and Illumina adapters (Bioo Scientific, Austin, TX, USA) were then added using the Kapa Hyper Prep Kit (KapaBiosystems, Wilmington, MA, USA). Ligation products were purified with AMPure XP beads (Beckmann Coulter Genomics, Danvers, MA, USA). The library was then quantified by qPCR using the KAPA Library Quantification Kit for Illumina Libraries (KapaBiosystems), and library profile was assessed using a High Sensitivity DNA kit on an Agilent Bioanalyzer (Agilent Technologies, Santa Clara, CA, USA). The library was sequenced on an Illumina NovaSeq instrument (Illumina, San Diego, CA, USA) using 150 base-length read chemistry in paired-end mode.

For Oxford Nanopore Technologies (ONT) sequencing, the library was prepared using the 1D Native barcoding genomic DNA (with EXP-NBD104 and SQK-LSK109). Genomic DNA fragments (1 µg) were repaired and 3’-adenylated with the NEBNext FFPE DNA Repair Mix and the NEBNext® Ultra™ II End Repair/dA-Tailing Module (New England Biolabs, Ipswich, MA, USA). Adapters with barcode provided by Oxford Nanopore Technologies (Oxford Nanopore Technologies Ltd, Oxford, UK) were then ligated using the NEB Blunt/TA Ligase Master Mix (NEB). After purification with AMPure XP beads (Beckmann Coulter, Brea, CA, USA), the sequencing adapters (ONT) were added using the NEBNext Quick T4 DNA ligase (NEB). The library was purified with AMPure XP beads (Beckmann Coulter), then mixed with the Sequencing Buffer (ONT) and the Loading Bead (ONT) and loaded on a MinION R9.4.1 flow cell. Reads were basecalled using Guppy 3.1.5.

### RNA extraction, library preparation and sequencing

When the cell concentration reached 10 million cells/mL in mid-exponential growth phase, 160 mL of culture were collected by three successive filtrations on 1.2 µm polycarbonate filters of 47mm to avoid prokaryotic contamination. To preserve cell and RNA integrity, we kept filtration time and pressure below 10 min and 20 mmHg, respectively. Then filters were stored in 15 mL Falcon tubes with 3 mL of Trizol (Invitrogen, Carlsbad, CA, USA), mixed and flash-frozen in liquid nitrogen for further processing. RNA was extracted by incubation at 65°C for 15 min, followed by a chloroform extraction. Aqueous phase was purified using Purelink RNA Isolation kit (Ambion Invitrogen, Carlsbad, CA, USA) according to the manufacturer’s instructions. DNA contamination was removed by digestion using the TURBO DNA-free™ Kit (Ambion Invitrogen) according to the manufacturer’s DNase treatment protocol. After two rounds of 30 min incubation at 37 °C, the efficiency of DNase treatment was assessed by PCR. Quantity and quality of extracted RNA were analyzed with RNA-specific fluorimetric quantitation on a Qubit 2.0 Fluorometer using Qubit RNA HS Assay (Invitrogen). The qualities of total RNA were checked by capillary electrophoresis on an Agilent Bioanalyzer, using the RNA 6,000 Nano LabChip kit (Agilent Technologies, Santa Clara, CA).

RNA-Seq library preparation was carried out from 1 µg total RNA using the TruSeq Stranded mRNA kit (Illumina, San Diego, CA, USA), which allows mRNA strand orientation. Briefly, poly(A)+ RNAs were selected with oligo(dT) beads, chemically fragmented and converted into single-stranded cDNA using random hexamer priming. After second strand synthesis, double-stranded cDNA was 3’-adenylated and ligated to Illumina adapters. Ligation products were PCR-amplified following the manufacturer’s recommendations. Finally, ready-to-sequence Illumina library was quantified by qPCR using the KAPA Library Quantification Kit for Illumina libraries (KapaBiosystems, Wilmington, MA, USA), and evaluated with an Agilent 2100 Bioanalyzer (Agilent Technologies, Santa Clara, CA, USA). The library was sequenced using 101 bp paired end read chemistry on a HiSeq2000 Illumina sequencer. Low-quality nucleotides (Q < 20) from both ends of the reads were discarded. Illumina sequencing adapters and primer sequences were removed and reads shorter than 30 nucleotides after trimming were discarded.

These trimming and cleaning steps were achieved using in-house-designed software based on the FastX package (https://www.genoscope.cns.fr/externe/fastxtend/). The last step identifies and discards read pairs that are mapped to the phage phiX genome, using the SOAP aligner^26^ and the Enterobacteria phage PhiX174 reference sequence (GenBank: NC_001422.1). This processing, described in Alberti et al, resulted in high-quality data^27^. Moreover, ribosomal RNA-like reads were excluded using SortMeRNA^28^.

### Long-read based genome assembly

Raw nanopore reads were used for genome assembly. Taxonomic assignation was performed using Centrifuge 1.0.3 to detect potential contamination. Genome size and heterozygosity rate were estimated using Genomescope^29^ and Illumina short-reads. For the genome assembly, we generated three sets of ONT reads: all the reads, 30x genome coverage with the longest reads and 30x genome coverage of the highest-scored reads estimated by Filtlong tool (https://github.com/rrwick/Filtlong). We then applied four different assemblers, Smartdenovo, Redbean, Flye and Ra on these three sets of reads (Table S1)^30–32^. After the assembly phase, we selected the best assembly (Flye with all reads) based on the cumulative size and contiguity. The assembler output was polished three times using Racon with Nanopore reads, and two times with Hapo-G and Illumina reads^33,34^. Gene completeness of the assembly was estimated using the single-copy orthologous gene analysis from BUSCO v5 with the stramenopile dataset version 10 containing 100 genes^35^.

### Repeat masking and GC analyses

Repetitive regions on the genome were masked using Tandem Repeat Finder tool^36^, Dust tool to detect low complexity regions^37^ and RepeatMasker^38^ to identify interspersed repeats based on homology search within the Stramenopile clade and other low complexity sequences. The positions of detected repeats were merged and hard-masked on the genome, amounting to a total of 8% of its length. *Ab initio* identification of repeat family sequences was performed using RepeatScout^39^. The algorithm first calculates the frequency of all k-mers in the genome, then removes low-complexity regions and tandem repeats. In > 80% of the cases, repeat families identified using *ab initio* approaches do not overlap with repetitive regions identified by homology search. GC content along the genome was calculated with Bedtools nuc version 2.29.2^40^ and the coverage over a non-overlapping window of 2 Kb with Mosdepth version 0.2.8^41^.

### Transcriptome assembly

RNA sequencing reads from *P. calceolata* RCC100 were assembled using Velvet 1.2.07 and Oases 0.2.08 with a k-mer size of 63 bp^42,43^. Reads were mapped back to the contigs with BWA-mem^44^ and only consistent paired-end reads were kept. Uncovered regions were detected and used to identify chimeric contigs. In addition, open reading frames (ORF) and domains were searched using respectively TransDecoder (http://transdecoder.sourceforge.net) and CDDsearch^45^. Contig extremities without predicted ORFs or functional domains were removed. Lastly, we used the read strand information to correctly orient RNA contigs. We completed the RNA contigs dataset with the two transcriptome assemblies of the RCC100 strain of *P. calceolata* from the Marine Eukaryotes Transcriptomes database (METdb) (http://metdb.sb-roscoff.fr/metdb/)^46^.

### Gene prediction

Nuclear gene prediction was performed using 23,696 Pelagomonadales proteins (mainly *Aureococcus anophagefferens*) downloaded from the NCBI website. Proteins were aligned on the genome in a two-step strategy. First, BLAT (version 36 with default parameters) was used to rapidly localize corresponding putative regions of these proteins on the genome. The best match and the matches with a score greater than or equal to 90% of the best match score were retained. Then, the regions with BLAT alignments were masked and we aligned the same set of proteins using BLAST, which is able to identify more divergent matches. Second, alignments were refined using Genewise (version 2.2.0 default parameters, except the -splice model option to detect non-canonical splicing sites), which is more accurate for detecting intron/exon boundaries. Alignments were kept if more than 50% of the length of the protein is aligned on the genome. Additionally, the transcriptome assemblies of *P. calceolata* RCC969, RCC2362, RCC706 and RCC981 included in the METdb were translated into proteins and aligned to the genome using BLAT, a BLAT score > 50 % filter, and alignments refined with Genewise as previously described.

We selected alignments from the newly generated transcriptome assembly and the two assemblies available in METdb belonging to the *P. calceolata* CCMP1214 strain to build a training set for AUGUSTUS *ab initio* gene predictor. Only gene models with complete coding DNA sequences were retained for training and 1,000 genes were set aside for testing AUGUSTUS accuracy. Initial training produced exon and intron parameters for *P. calceolata* species. Parameters were optimized using successive steps of training and testing. We calculated the accuracy of gene prediction by running AUGUSTUS on the test set. At the exon level, AUGUSTUS performed well in terms of sensitivity (0.619) and specificity (0.669). We thus run AUGUSTUS on the masked genome based on trained parameters. The *ab initio* prediction and all the transcriptomic and protein alignments were combined using Gmove which is an easy-to-use predictor with no need for a pre-calibration step^47^. Briefly, putative exons and introns, extracted from predictions and alignments, were used to build a graph, where nodes and edges represent exons and introns respectively. From this graph, Gmove extracts all paths and searches open reading frames (ORFs) which are consistent with the protein evidence. We trimmed untranslated transcribed regions that overlap the coding part of a neighbour gene and renamed the genes following the standard nomenclature. Mono-exonic genes models encoding proteins of less than 200 amino acids without significant protein match (1,006 genes) were excluded. Chloroplast and mitochondrial genes (contig 7 and 8) were predicted using previously published annotations for *P*.*calceolata*^22,48^. Following this pipeline, we predicted 16,667 genes with 1.45 exons per gene on average.

The presence of introner elements (IE) in *P. calceolata* was investigated by overlapping the position of the introns in transcriptome alignments with the positions of repeat families detected *ab initio* with RepeatScout^39^. Intron overlapping repeats over more than 90% of their length were identified as putative IEs.

### Functional analysis

Predicted gene models of *P. calceolata* nuclear genome (contig 1 to contig 6) were annotated for protein function using InterProScan v5.41-78.0^49^. A protein alignment against the NR database (01-12-2021 version) was performed with diamond v0.9.24^50^. The best protein match with a functional annotation and an e-value < 10^−5^ was retained. KEGG Orthologues (KO) were identified with the HMM search tool KofamKoala v1.3.0 and KO annotations with an e-value <10^−5^ and a score above the HMM threshold were retained^51^. Finally, Gene Ontology (GO) terms and Enzyme commission (EC) numbers were recovered from the interproscan and KO analysis respectively. Previously published chloroplast and mitochondrial gene names and functions were reported on the corresponding genes^22,48^. All gene functional annotations of *P. calceolata* are available in Table S2.

In order to compare the functional annotation of *P. calceolata* with other small free-living photosynthetic eukaryote, we applied the same analysis on the predicted proteins available for the following species: *Aureococcus anophagefferens, Thalassiosira pseudonana, Phaeodactylum tricornutum, Nannochloropsis oceanica, Bathycoccus prasinos, Micromonas pusilla, Ostreococcus lucimarinus* and *Emiliania huxleyi* (references are indicated in Table 1).

**Table 1:** Nuclear genome characteristics of several unicellular photosynthetic eukaryotes.

| Phylum/Class | Species | Genome size (Mb) | Number of chromosomes | Predicted genes | GC% | Cell size | References |
| --- | --- | --- | --- | --- | --- | --- | --- |
| Pelagophyceae | *Pelagomonas calceolata* | 32.4 | 6 | 16,613 | 63.6 | 2 µm | This study |
| Pelagophyceae | *Aureococcus anophagefferens* | 56.0 | Unkown | 11,520 | 67.4 | 2 µm | 66 |
| Eustigmatophyceae | *Nannochloropsis oceanica* | 29.3 | 32 | 7,730 | 54.0 | 3 µm | 67 |
| Diatom | *Phaeodactylum tricornutum* | 27.0 | 33 | 10,402 | 48.8 | 11 µm | 68 |
| Diatom | *Thalassiosira pseudonana* | 34.5 | 24 | 11,776 | 46.9 | 5 µm | 69 |
| Chlorophyta | *Micromonas pusilla* | 21.9 | 17 | 10,575 | 65 | ≤2 µm | 70 |
| Chlorophyta | *Ostreococcus lucimarinus* | 13.2 | 21 | 7,651 | 60 | 1.3 µm | 71 |
| Chlorophyta | *Bathycoccus prasinos* | 15.1 | 19 | 7,847 | 48 | 1-2 µm | 72 |
| Haptophyta | *Emiliania huxleyi* | 141.7 | Unkown | 38,549 | 64.5 | 4-5 µm | 73 |

We defined a list of 23 meiosis-specific genes using three previous studies^52–54^. KO annotations and Interproscan domains were used to recover these genes in the *P. calceolata* genome.

NIT-sensing domain containing genes were identified based on Interproscan annotations. Eukaryotic homologs of the 3 *P. calceolata* NIT-sensing genes were retrieved with a BLASTP (e-value < 10^−5^, coverage > 100 aa) against 27.7 million proteins from NR, the METdb^46^ transcriptome database, *Tara* Oceans single-cell amplified genomes and metagenome assembled genomes (SMAGs)^55^ and eukaryotic algal proteomes from the JGI database. The 60 retrieved proteins and the 3 *P. calceolata* NIT-domain containing proteins were then aligned with Mafft v7.0 (https://mafft.cbrc.jp/alignment/server/) and a Maximum Likelihood phylogenetic tree (Jones-Taylor-Thornton substitution model and 100 bootstraps) was made with MEGAX software. Transmembrane regions in NIT-domain containing proteins were identified with TMHMM v 2.055.

### Mapping and filtering of environmental metagenomic reads

We used metagenomics datasets of *Tara* Oceans and *Tara Polar Circle* expeditions to detect the *P. calceolata* genome in the oceans. Datasets from water samples collected on the photic zone: surface (SUR) and deep-chlorophyll maximum (DCM) were analysed. Size-fractionated water samples containing pico-nano algae (organisms <5µm in cell diameter) were selected: 0.2-3 µm (100 samples), 0.8-2000 µm (119 samples) and 0.8-5 µm (148 samples)^27^. Metagenomic reads were aligned on the *Pelagomonas calceolata* genome with BWA-mem version 0.7.15 with default parameters^56^. Alignments with at least 90% of identity over 80% of read length were retained for further analysis. In the case of several possible best matches, a random one was picked. In order to remove putative PCR duplicates, multiple read pairs aligned at the exact same position on the *P. calceolata* genome were removed with samtools rmdup version 1.10.2^44^.

### Relative abundance calculation

The relative abundance of *P. calceolata* was calculated from both metabarcoding data and metagenomic data. For the metabarcoding abundance, we used the 18SV9 rRNA OTU table published in 2021 and available here https://zenodo.org/record/3768510#.YEX2S9zjJaQ^57^. Bacterial and archaea OTUs were removed for this analysis. For the metagenomic abundance, we divided the total number of reads aligned on the *P. calceolata* genome by the total number of sequenced reads for each sample.

### *P. calceolata* relative abundance modelling

Two *P. calceolata* relative abundance models were performed based on the *in situ* environmental conditions measured at the time of sampling or using the World Ocean Atlas 2018 (WOA18) datasets. The environmental parameters measured during the expedition are available in the Pangaea database (https://www.pangaea.de/) and are described in^58^. Iron concentrations are annual means derived from PISCES2 model^59^ and described in^12^. Ammonium concentrations at the date and location of sampling are derived from the MITgcm Darwin model and available in the Pangaea database^60^. Environmental parameters for each sample are available in Table S3. Pearson’s correlation between the relative abundance of *P. calceolata* and all environmental parameters were calculated with the *cor* function in R and the GGally package version 2.1.0. The 8 parameters with a correlation above 0.2 or below -0.2 were selected for downstream analysis. Principal component analysis (PCA) was performed with the R package FactoMineR version 2.4. We used a Generalized Additive Model (GAM) for its ability to fit non-linear and non-monotonic functions and for its low sensitivity to extreme values to model the relative abundance of *P. calceolata* as a function of iron concentration and temperature^61^. This function is implemented in the mgcv R package version 1.8-33. All figures were generated with R version 4.0.3 and ggplot2 package version 3.3.2.

The mean of several climatologies of the Earth System Models under RCP8.5 scenario (ESM8.5) were used to define the environmental conditions at the end of the century following the method of Frémont et al^62^. *P. calceolata* relative abundance modelling based on the World Ocean Atlas 2018 (WOA18) at locations, depths and months of *Tara* Oceans samples or ESM8.5 were obtained using four machine learning techniques^62^. Briefly, Generalized Additive Models (gam), Neural Networks (nn), Random Forest (rf) and Gradient Boosted Trees (bt) were trained in regression mode. Over 100-fold random cross validation, all models performed well: mean Pearson correlation coefficient were respectively 0.64 (gam), 0.67 (nn), 0.71 (rf) and 0.70 (bt). We used the ensemble model approaches^63^ for final global-scale modelling of the relative abundance of *P. calceolata* (i.e. the mean projections of the validated machine learning techniques).

## Results

### A compact genome revealed by a chromosome-scale assembly

To investigate its gene repertoire and its distribution across the oceans we sequenced and assembled the genome of *P. calceolata* using ONT long-reads and Illumina short-reads. Using k-mer distribution of short reads, the genome was estimated to be homozygous with a size of 31 Mb (Figure S1). The ONT long-reads were assembled using Flye into 6 nuclear contigs for a total of 32.4 Mb, 1 plastid circular contig (90 Kb) and 1 mitochondrial circular contig (39 Kb) (Figure 1 and Figure S2A,B). Two large and highly similar duplicated regions (>99% of identity) were detected at the extremity of contig 1 and 5 (393 Kb) and at the extremities of contig 3 and 6 (192 Kb). The vertical read coverage of these two regions is similar to that of other genomic regions suggesting that they are duplicated in all cells of this *P. calceolata* culture (Figure 1 and Figure S2C). In addition, 150 Kb at one extremity of contig 4 present a higher vertical coverage suggesting that this region is also duplicated in the *P. calceolata* genome but collapsed in the assembly (Figure S2). Interestingly, duplicate regions at the end of sequences have already been observed in the chlorophyte *Ostreococcus tauri* which has been maintained for several years in culture^64^. These observations suggest that culture could affect not only the sequence of a given organism’s genome but also its structure. (TTAGGG)n telomeric repeats were detected at both ends of contigs 2, 3, 4 and 6 indicating that these 4 contigs represent complete chromosomes (Figure S2). For contig 1 and contig 5, telomeric sequences are present at only one extremity, the other extremity ending in the duplicated region. This result suggests that the six contigs correspond to six chromosomes of *P. calceolata*.

**Figure 1:**
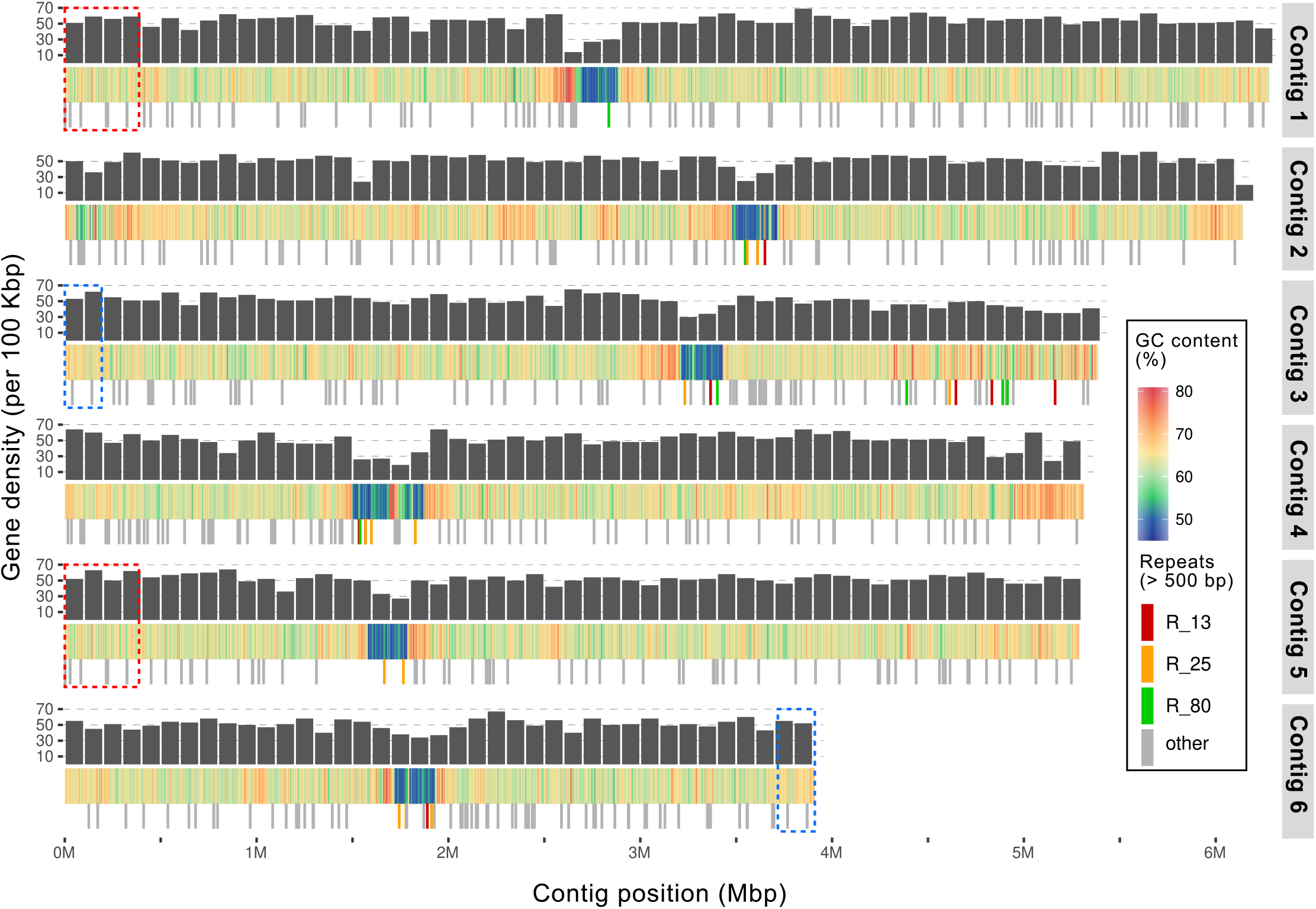
*Pelagomonas calceolata* nuclear genome. Representation of the 6 nuclear contigs of *P. calceolata*. The top layer indicates the number of genes per 100 Kb (black bars), the middle layer represents the GC content in percentage over a window of 200 Kb and the bottom layer is the position of DNA repeats of more than 500 bases repeated at least 5 times over the entire genome. Red, orange and yellow bars indicate three different repeats in low-GC regions present in at least 3 different contigs. Dashed red and blue rectangles are duplicated chromosomic regions.

A total of 16,667 genes was predicted on the *P. calceolata* genome with an average of 1.45 intron per gene (Table 1). The distribution of intron lengths reveals a peak at around 210 bp (Figure S3A), which is the characteristic length of Introner Elements (IE) described in *A. anophagefferens*^65^. Putative IE were then identified (n=956, see method). Both canonical (GT) and non-canonical (GC) donor splicing sites were present at the ends of putative IE, while the acceptor site was canonical (AG) in all cases (Table S4). The logo representation of the putative IE sequences reveals the presence of GT and GC donor sites at the 5’ end, AG acceptor site at the 3’ end, and conserved TIR in the flanking regions (Figure S3B). 9, 583 *P. calceolata* predicted proteins (58%) are homologous with at least one gene in a Stramenopile genome, including 2,631 (16%) only shared with the Pelagophyceae *A. anophagefferens* (Figure S4). A conserved functional domain (Pfam, KO or InterProScan) was found in 11,240 proteins (67%).

### GC content distribution along *P. calceolata* chromosomes

A remarkable feature in *P. calceolata* genome is the distribution of GC content along *P. calceolata* chromosomes (Figure 1). While the average GC content of the nuclear genome is 63%, one large region in each contig (259 Kb in average) is 52% GC. These unique large troughs in GC content in each chromosome suggest that these regions may encompass centromeres. We did not observe accumulation of repeated elements or transposons in these low-GC regions; however, the gene structures differ from that of other genomic regions (Figure 1 and Table S5). The slight decrease of gene density observed in Figure 1 is explained by a longer gene size (average of 3 Kb compared to 1.9 Kb in other regions). In addition, the 453 genes in low-GC regions contain more introns (2.56 introns per gene compared to 0.49) and the introns are shorter (120 compared to 214 bases). We also noticed a higher proportion of bases belonging to intergenic regions, 18% in low-GC regions and 11% in other chromosomic regions (Table S5). Interestingly, three repeats of more than 500 bases are present in several low-GC regions (R_13 in contigs 2, 3 and 6; R_25 in all contigs except contig 1 and R_80 in contigs 1 to 4) (Figure 1). No homology was found between these sequences and known repeat elements.

Gene function analysis of low-GC regions reveals an enrichment of genes involved in specific cellular mechanisms (Table S6). Thirteen genes are involved in DNA replication including the Anaphase-promoting complex subunit 4, the sister chromatid cohesion protein Dcc1 and 3 Mini-chromosome maintenance genes. Twenty-five genes are involved in microtubule synthesis and microtubule-binding motor proteins (9 genes carrying dynein domains, 6 genes carrying kinesin motor domains and 4 tubulin genes). These genes indicate that the low-GC regions contain many genes required for *P. calceolata* cellular division. Finally, 18 genes are involved in transcription including 3 genes encoding RNA Pol II rpb2 subunits and 7 genes encoding transcription factors suggesting an important role of these chromosomic regions for the regulation of gene expression.

### Sex related genes in *P. calceolata*

Because low-GC regions could be related to meiotic recombination^74^, we looked for genes involved in sexual reproduction in the *P. calceolata* genome. Among 20 genes specifically involved in meiosis, 16 homologs are present in the *P. calceolata* genome (Table 2). These genes include the double-strand DNA break (DSB) initiator SPO11; RAD50, RAD52 and MRE11 to bind DSBs; HOP2, MND1, DMC1 and RAD51 to ensure pairing and invade the homologous strand; MSH2, MSH3 PMS1 and MSH6 genes involved in the synthesis-dependent strand annealing pathway and MUS81 necessary for non-interfering (class II) crossing over. Interestingly, MSH4 and MSH5 genes are absent from the *P. calceolata* genome. Indeed, these genes necessary to perform the interfering (class I) recombination pathway through Double Holliday Junctions are present in most eukaryotic lineages. The large low-GC regions could be a consequence of the absence of the MSH4/5 genes as suggested for yeasts^75^ (see discussion). There are also no homologs of ZIP, HOP1 and RED1 genes in the *P. calceolata* genome. These genes are known to be involved in homologous pairing of chromatids and construction of the synaptonemal complex in animals, plants and fungi but are not required to perform meiosis. Indeed these 3 genes are absent in several phyla capable of meiosis like diatoms^76^ and ciliates^52^. Taken together, the genetic content of *P. calceolata* strongly suggests that this species perform meiosis.

**Table 2:**
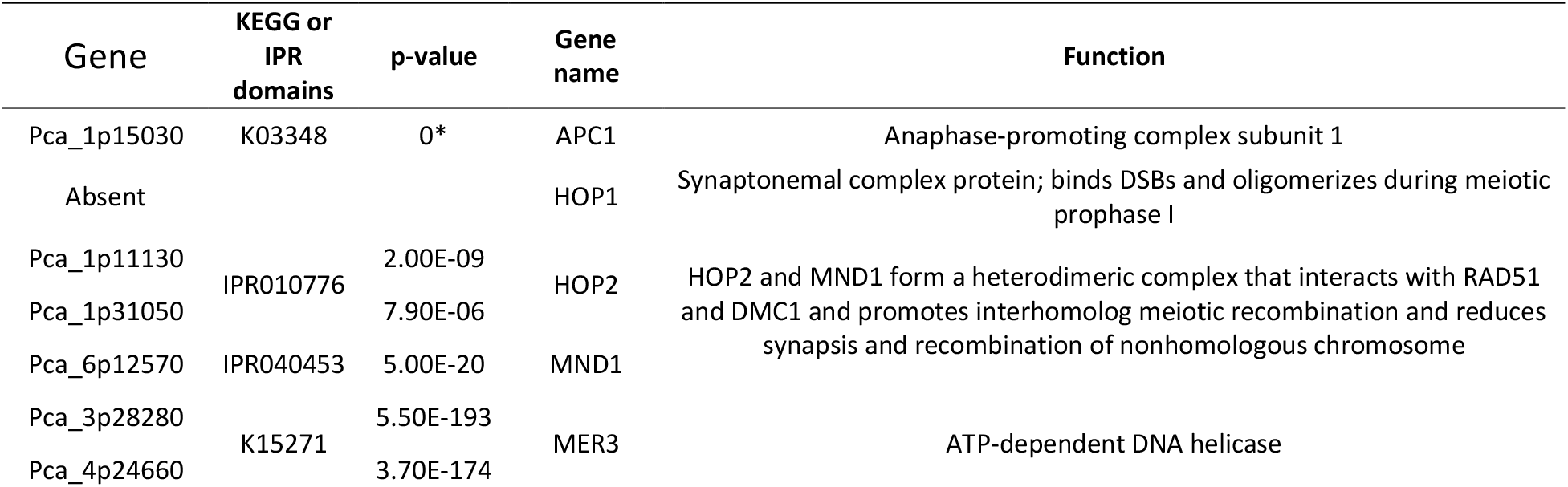
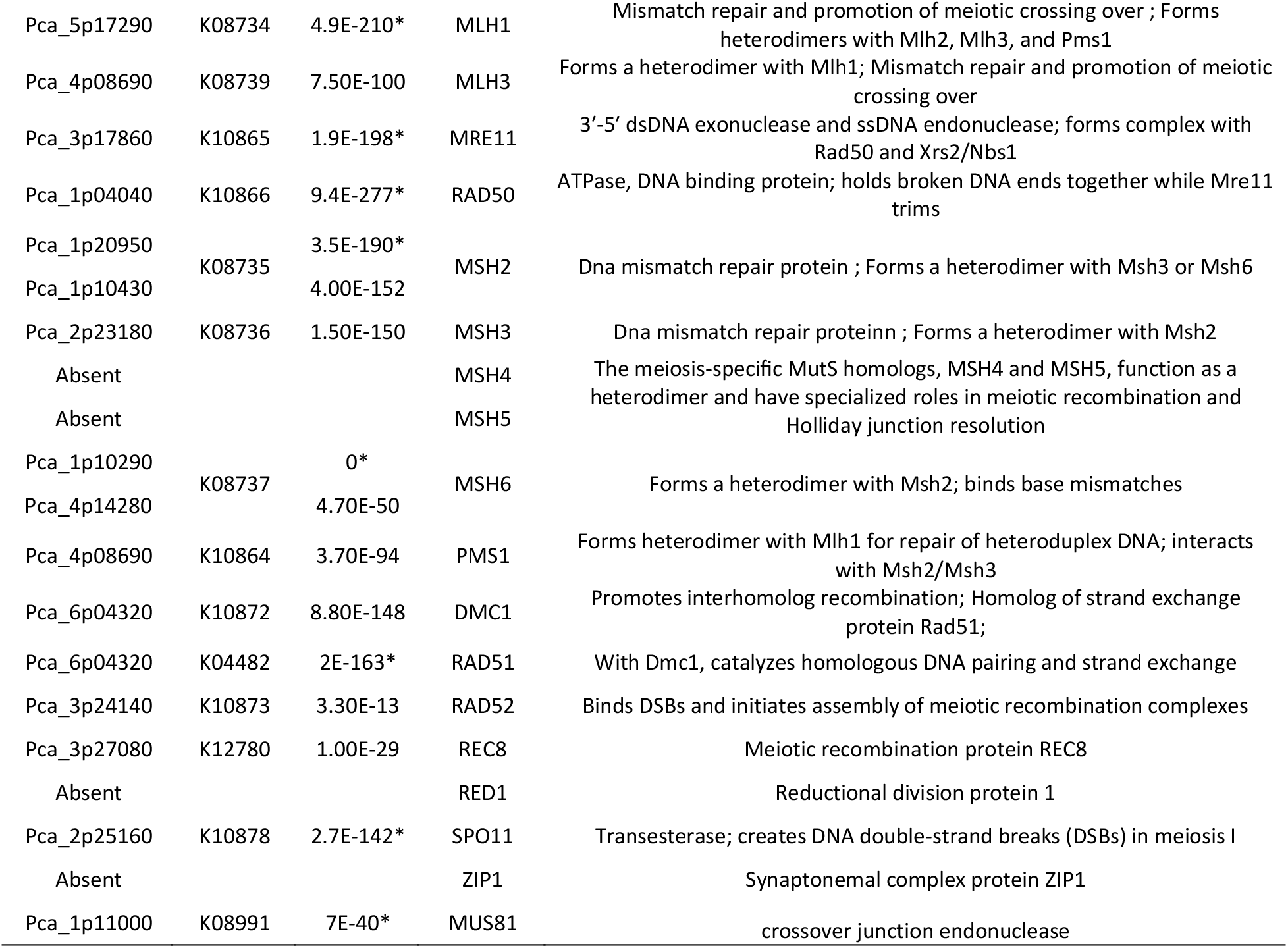
Putative meiotic genes identified in *P. calceolata* genome. Kegg Orthology p-values with a star are above the HMM threshold defined by KoFamKoala tool.

### Genes involved in nitrogen uptake, storage and recycling

*P. calceolata* could be an important player in the nitrogen (N) cycle in oceanic ecosystems^24^, therefore we explored the genomic capacities of *P. calceolata* to assimilate and use nitrogen-containing compounds. We systematically identified and counted genes involved in the nitrogen cycle in *P. calceolata* compared to seven other pico-nano photosynthetic eukaryotes (Figure 2 and Table S7). The uptake of nitrogen-containing inorganic compounds is supported by 13 genes in *P. calceolata*. Among them, 8 genes encode nitrate/nitrite or formate/nitrite transporters, which is on average higher than in other algae. In contrast, only 5 genes encode ammonium transporters which is low compared to other species suggesting that nitrite and/or nitrate is the main external source of inorganic nitrogen for *P. calceolata*. The number of enzymes incorporating ammonium into organic compounds (GS/GOGAT pathway) are higher in *P. calceolata* than in other species: 5 glutamine synthetase and 4 glutamate synthase genes are present in the *P. calceolata* genome. On the contrary, fewer genes assure the reduction of nitrate into nitrite (1 nitrate reductase) then nitrite into ammonium (2 nitrite reductases) (Table S7).

**Figure 2:**
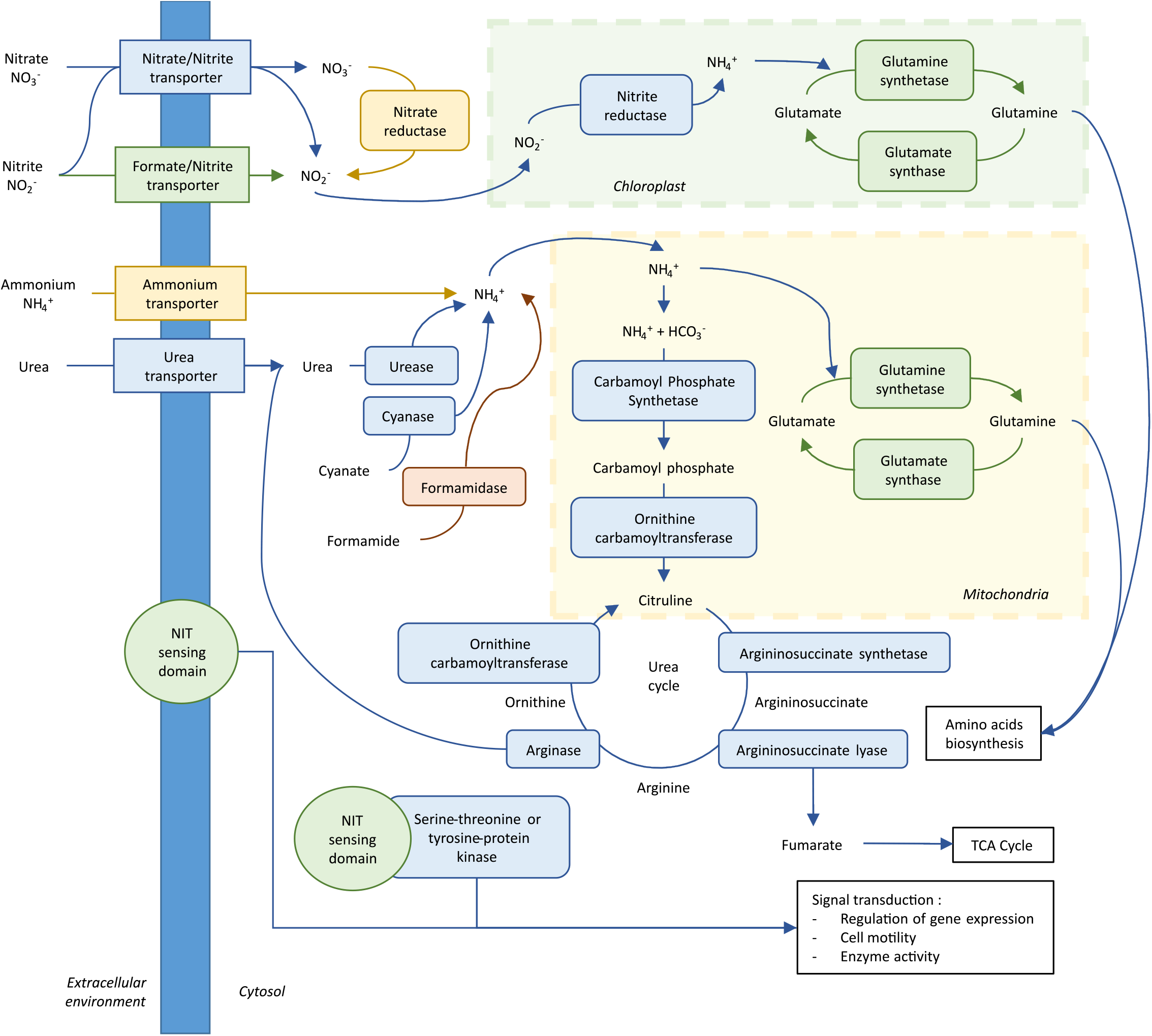
Schematic representation of N transport and assimilation in *P. calceolata* based on the gene content. Rectangles represent transporters of inorganic nitrogenous compounds, ovals represent enzymes and chemicals are unboxed text. The color code indicates if the number of gene copies for a specific function is overrepresented (green), equally represented (blue), underrepresented (orange) or absent (red) in *P. calceolata* compared to the mean of 8 pico-nano photosynthetic eukaryotes. Gene copy number for each function is indicated in Table S7.

We identified 3 genes carrying the nitrate and nitrite sensing (NIT) domain (IPR013587) in the *P. calceolata* genome. One *P. calceolata* protein carries an NIT-sensing domain surrounded by 2 transmembrane domains suggesting a capacity for external Nitrate/Nitrite sensing while the 2 other NIT genes carry a protein-kinase domain (IPR000719) suggesting a phosphorylation-based signal transduction dependent on intracellular nitrate or nitrite concentration (Figure S5). Using NCBI non-redundant proteins and marine genomic databases (see methods), 60 homologous proteins of the NIT-sensing domains of *P. calceolata* were identified. These homologs are restricted to the Pelagophyceae class (16 transcriptomes), the Dictyochophyceae class (6 transcriptomes) and one putative Cryptophyte transcriptome. The phylogenetic tree of this protein family shows three subfamilies diverging before the Dictyochophyte/Pelagophyte separation (Figure S5).

We then identified genes involved in nitrogen recycling from organic compounds which are important in several species in case of inorganic nitrogen deprivation. One arginase gene and one cyanase gene were detected in the *P. calceolata* genome but no gene encoding formamidase. In addition, the number of gene copies for enzymes involved in the urea cycle (carbamoyl-phosphate synthetase, ornithine carbamoyltransferase, argininosuccinate synthase and argininosuccinate lyase) is equal or slightly lower than in other algae (Figure 2 and Table S7). These results suggest that *P. calceolata* is not particularly adapted to recycle nitrogen from organic molecules but could be capable of incorporating inorganic nitrogen compounds even in N-poor environments.

### Genes related to iron uptake, storage and usage in *P. calceolata*

Iron is a critical metal for all photosynthetic organisms. Iron-containing molecules are essential for the photosynthesis, the nitrogen cycle and the protection against reactive oxygen species. We identified gene coding for iron uptake and storage in *P. calceolata* genome and compared them to other small photosynthetic eukaryotes (Table S7). *P. calceolata* has 5 genes encoding the phytotransferrin ISIP2 involved in Fe^3+^ uptake via endosomal vesicles and 2 putative iron storage protein ISIP3. These genes are transcribed following starvation in diatoms suggesting a capability for thriving in iron-poor environments for *P. calceolata*. In addition, 3 genes encode the iron transporter ferroportin. These proteins are iron exporters in multicellular organisms but their function in micro-algae remains to be studied^77^. Zinc/iron permeases, transmembrane ferric reductases and multicopper oxidase genes, involved in iron uptake from the environment, are present in the *P. calceolata* genome (8, 5 and 2 genes respectively) but in equal or lower gene copy number than in other studied species. The iron permease FRT1 and the ISIP1 gene, involved in endocytosis of iron-chelator molecules (siderophores), are absent in the *P. calceolata* genome. Furthermore, *P. calceolata* has no gene encoding the iron storage protein ferritin. The absence of these genes suggests that iron uptake and storage is not a major asset of *P. calceolata* compared to the other photosynthetic protists.

Several important ferrous proteins can be substituted by non-ferrous equivalents in iron-poor environments^13^. In the *P. calceolata* genome we identified 11 flavodoxin and 3 phytocyanin genes, encoding non-ferrous proteins involved in electron transfer during photosynthesis potentially replacing Ferredoxin and Cytochrome C6 respectively. The Fructose-bisphosphate Aldolase (FBA) necessary for gluconeogenesis and the Calvin cycle is encoded by 6 genes in *P. calceolata*. Two genes are dependent on a bivalent cation (FBA type II), the four others are Zinc/Iron-independent (FBA type I). Finally, all types of Superoxide dismutases (SOD) are found in the *P. calceolata* genome. Non-ferrous SODs (Cu/Zn and Ni) are encoded by 3 genes, while the Mn/Fe-SOD is encoded by 2 genes. The genetic content of *P. calceolata* indicates that several essential processes are potentially iron-independent thanks to the presence of many iron-free alternative proteins.

### Relative abundance of *P. calceolata* across oceanic basins

First, we used the OTU table computed from metabarcoding samples of the *Tara* Oceans expedition to estimate the relative abundance of *P. calceolata* across all oceans^57^. The relative rRNA abundance of *P. calceolata* OTU in the 0.8-5 µm size-fraction is 1.30% on average for the 111 surface samples and 0.81% on average for the 62 DCM samples (Table S8). According to this method of abundance estimation, *P. calceolata* is the third most abundant eukaryote of the 0.8-5 µm size-fraction among *Tara* samples covering all oceans except the Arctic Ocean. However, the number of rRNA copies in each organism biases this metabarcoding-based abundance estimation. Therefore, we used the mapping of metagenomic reads on the *P. calceolata* genome to estimate more precisely its abundance. The relative abundance obtained with the entire genome is strongly correlated to the metabarcoding-based relative abundance (Pearson correlations of 0.91 and 0.70 for 0.8-2000 and 0.8-5 µm size-fractions respectively). However, the metabarcoding-based abundance is underestimated by a factor of 2.3 in the 0.8-5 µm size-fraction and a factor of 3.1 in the 0.8-2000 size-fraction compared to the metagenomic-based abundance (Figure S6). This underestimation is probably due to the low copy number of rRNA in *P. calceolata* (2 complete copies) compared to most other species with larger genomes.

The relative abundance of *P. calceolata* calculated with the number of reads aligned on the genome is estimated above 1% in 66 oceanic stations with a maximum of 6.7% for the 0.8-5 µm size-fraction at the DCM in the North Indian Ocean (Figure 3A and Figure S7). In the Indian Ocean, Red Sea and Mediterranean Sea, *P. calceolata* is significantly more abundant in the DCM than at the surface (Figure 3B). In cold waters (below 10°C), *P. calceolata* is not detected above our threshold of 25% of genomic horizontal coverage. Important variations between and within each oceanic basin are observed, suggesting that many biotic or abiotic factors influence *P. calceolata* abundance.

**Figure 3:**
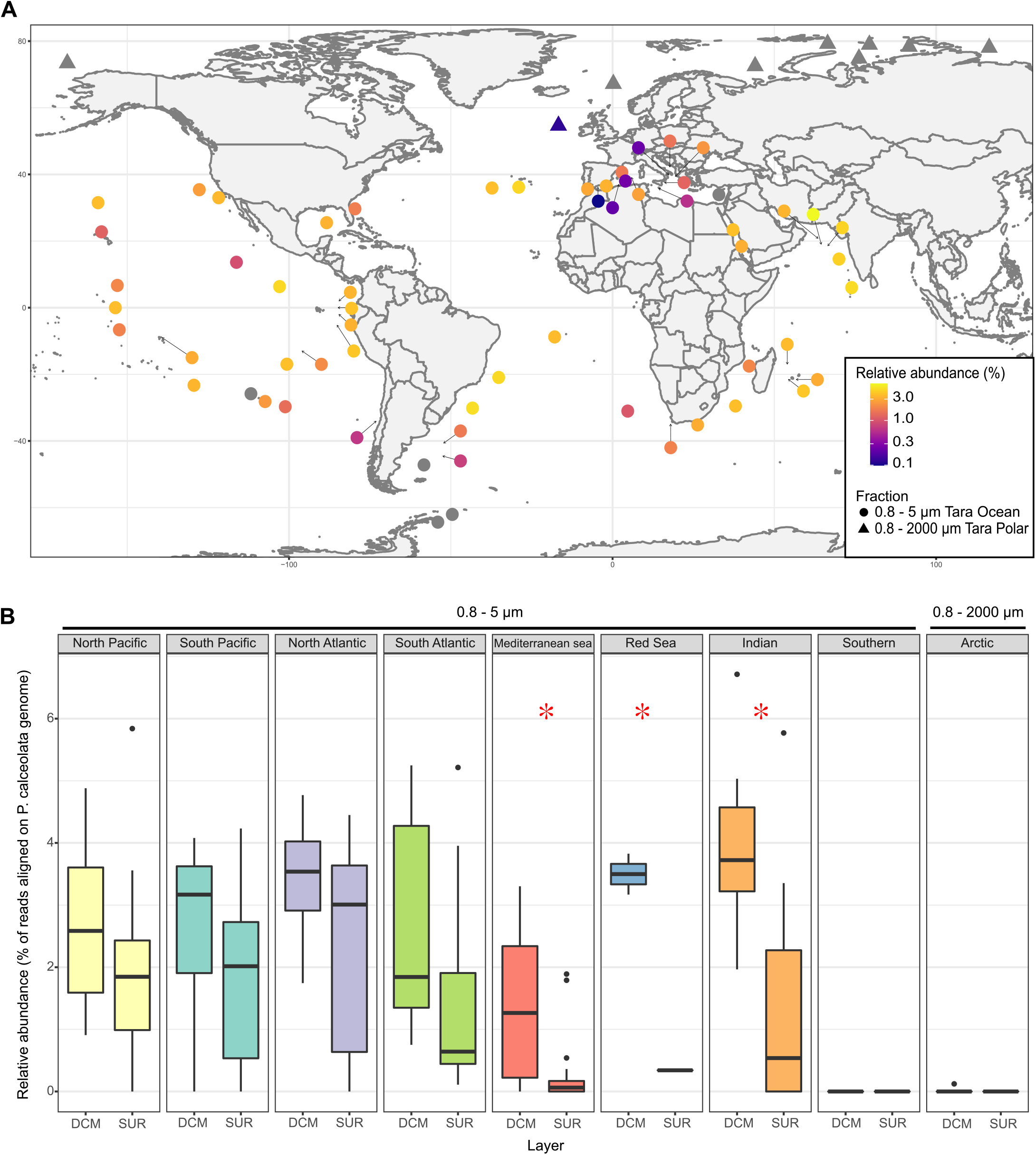
Relative abundance and distribution of *P. calceolata* in the oceans. A) World map of the relative abundance of *P. calceolata*. The colour code indicates the percentage of sequenced reads aligned on the *P. calceolata* genome. The DCM samples of size-fractions 0.8-5µm (circles) or 0.8-2000 µm (triangles) are shown. *P. calceolata* is considered to be absent if the horizontal coverage is below 25% of the genome (grey dots). B) Boxplot of the relative abundance of *P. calceolata* in each oceanic region in surface and DCM samples. Red stars indicate a significant difference between SUR and DCM samples (Wilcoxon test, p-value<0.01

### High relative abundance of *P. calceolata* in temperate and iron-poor regions

In order to identify factors controlling *P. calceolata* abundance in the oceans, we used physical-chemical parameters available for each oceanic station (see methods). Principal component analysis reveals a positive relation between *P. calceolata* abundance, the temperature and the coast distance and a negative relation with iron concentration (Figure 4A and B). The other parameters do not seem to be related to *P. calceolata* abundance. This result is consistent over the 3 size-fractions containing *P. calceolata* cells (0.8-5 µm, 0.8-2000 µm and 0.2-3 µm size-fractions) (Figure S8).

**Figure 4:**
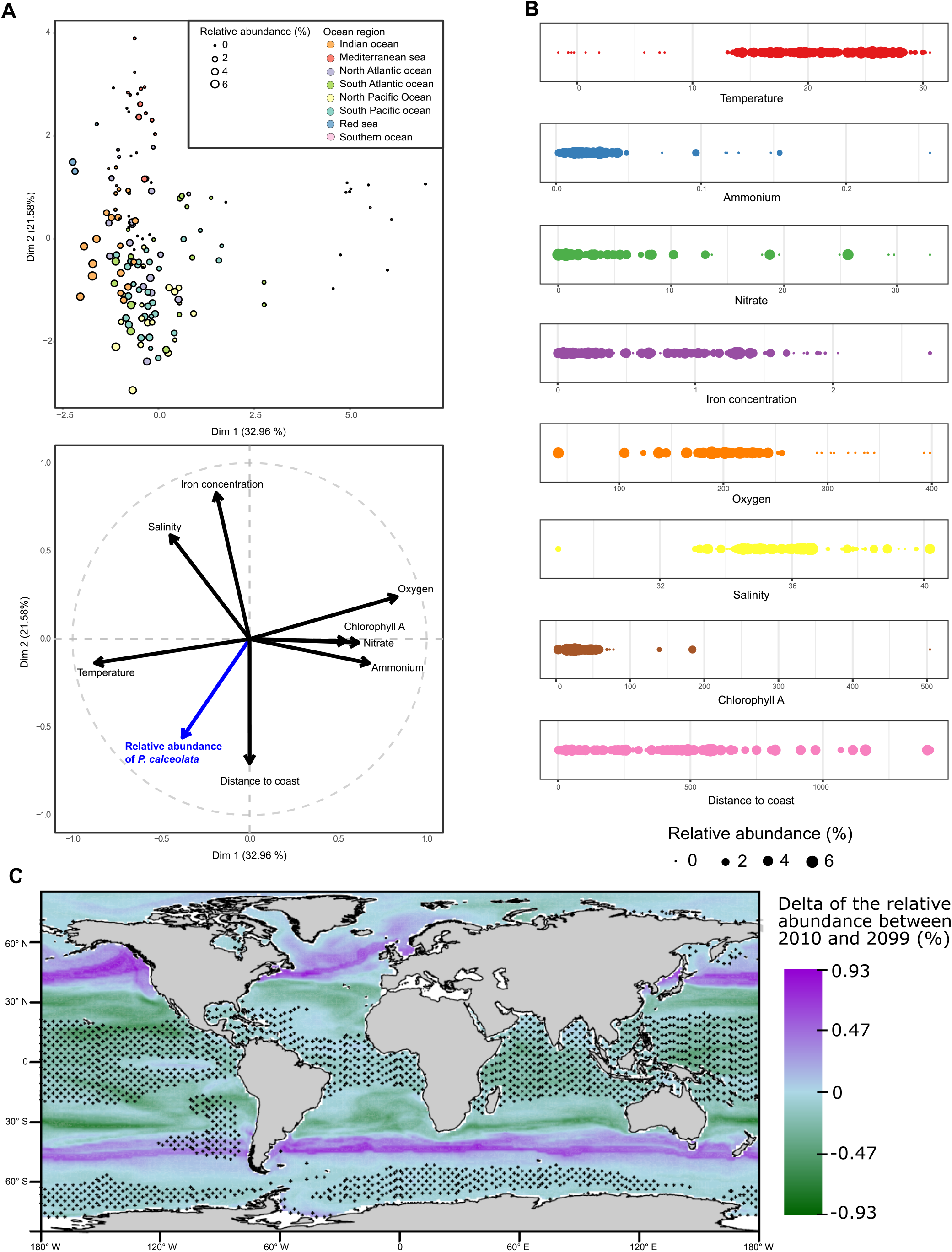
Ecological niche of *P. calceolata*. A) Principal component analysis of the relative abundance of *P. calceolata* and 8 environmental parameters measured during Tara sampling.Each dot represents a sample with a size proportional to the relative abundance of *P. calceolata*.The colours indicate the oceanic basins. B) Bubble plot of the relative abundance of P. calceolata for the 0.8-5 µm size-fraction according to the 8 environmental parameters. C) Difference of the modelled relative abundance of *P. calceolata* between 2010 and 2099. Green areas correspond to a decrease while purple areas correspond to an increase of *P. calceolata* relative abundance. Small stars indicate locations where at Ieast one of the predictor drivers is out of range compared to the training dataset values.

Despite the numerous factors potentially influencing *P. calceolata* abundance, we observed a weak but significant Pearson’s positive correlation with the temperature and a negative correlation with iron concentration (Table 3). In addition, we used a general additive model to estimate the contribution of the combination of temperature and iron concentration to *P. calceolata* relative abundance (Table 3). The two factors contribute significantly and explain 17.9% of the variations of *P. calceolata* abundance in the 0.8-5 µm size-fraction and 47.9% in the 0.8-2000 µm size-fraction. The high relative abundance of *P. calceolata* in iron-poor waters suggests that this species is particularly capable of acclimation to this environmental condition.

**Table 3:** Environmental parameters explaining *P. calceolata* relative abundance for the (A) 0.8-5 µm and the (B) 0.8-2000 µm size-fractions.

| A | 0.8 - 5 $\mu\text{m}$ | GAM model | | | GAM verification | | Pearson correlations | |
| --- | --- | --- | --- | --- | --- | --- | --- | --- |
|  |  | edf | F value | p-value | k-index | k p-value | r | p-value |
|  | s(Temperature) | 2.754 | 4.229 | 0.00506 | 1.00 | 0.46 | 0.23 | 0.001 |
|  | s(iron concentration) | 1.897 | 5.549 | 0.00326 | 0.92 | 0.16 | -0.25 | 0.001 |
|  | Adjusted R<sup>2</sup> | 0.151 |  |  |  |  |  |  |
|  | Deviance explained | 17.9% |  |  |  |  |  |  |
| B | 0.8 - 2000 $\mu\text{m}$ | GAM model | | | GAM verification | | Pearson correlations | |
|  |  | edf | F value | p-value | k-index | k p-value | r | p-value |
|  | s(Temperature) | 4.223 | 8.769 | 7.89e-07 | 0.98 | 0.405 | 0.57 | 0.0001 |
|  | s(iron concentration) | 1.450 | 0.793 | 0.348 | 0.86 | 0.045 | -0.47 | 0.0001 |
|  | Adjusted R<sup>2</sup> | 0.449 |  |  |  |  |  |  |
|  | Deviance explained | 47.9% |  |  |  |  |  |  |

Finally, we projected the relative abundance of *P. calceolata* at the end of the century. We modelled the ecological niche of *P. calceolata* using the WOA18 datasets at the time and location of sampling or using the projected climatology in 2099 using the RPC8.5 scenario (see methods). We projected an increase (from +0.4% to +0.92%) of *P. calceolata* relative abundance from latitude 40° to latitude 50° in the North and South hemispheres and a slight decrease in temperate and tropical waters (−0.5% maximum) (Figure 4C and Figure S9). The decrease in the tropical waters is uncertain (stars on the world map) because this environment in 2099 is out of the range of the training dataset.

## Discussion

The essential roles of phytoplankton in oceanic ecosystems have been illustrated many times, however numerous lineages are still poorly explored and model organisms are restricted to a few taxa (mainly diatoms, prasinophytes, and haptophytes) limiting the global understanding of phytoplankton activity. The *P. calceolata* genome assembled and annotated in this study brings new insights into specific genomic features of this algae class related to its adaptation to specific environments and reveals a previously underestimated high abundance of *P. calceolata* in the oceans.

### Large low-GC centromeres

The chromosome level assembly of the *P. calceolata* genome thanks to nanopore read sequencing revealed unique genomic features that are not usually studied in short-read assembled genomes. The most striking observation is the presence of a unique GC trough (50%) in each scaffold contrasting with the high GC content (63%) of other genomic regions of *P. calceolata*. In eukaryotes, centromeres have a large variety of structures and characteristics. They are composed of many repeated sequences or contain genes, they are determined genetically or epigenetically and their size can vary from 125 bases (in *S. cerevisiae*) to several Mb (in mammals)^78^. Short regional centromeres (1-5 Kb) are generally low-GC compared to the genome and the sequence is unique at each centromere. Among stramenopiles, centromeres were characterized in the diatom *P. tricornatum* where a short low-GC sequence (>500 bp) is enough to define a centromere^79^. Large regional centromeres (>10 Kb) are generally gene-free, contain repeated elements and are not transcribed^78^. *P. calceolata* putative centromeres derive from this general pattern with large low-GC regions containing genes carrying essential functions of DNA replication, transcription and microtubule assembly. Interestingly, the red alga *Cyanidioschyzon* as well as several yeast species seem to have similar centromere structures with large, low-GC centromeres containing genes^80,75^.

### GC content, meiosis and recombination

The main hypothesis to explain these low-GC patterns in centromeres is the importance of GC-biased gene conversions (gBGC) during recombination and the inhibition of this recombination near centromeres^81^. Indeed, gBGCs increase the GC content of recombining DNA over evolutionary time inducing GC content variations within and between genomes^74^. The kinetochore formation at centromeric regions inhibits recombination and double strand break formation during meiosis resulting in rare gBGC in these regions^82^. Centromeric and peri-centromeric regions may therefore have a lower GC content. Furthermore, the genetic content of *P. calceolata* indicates that this species is capable of meiosis despite the absence of some genes including MSH4 and MSH5 necessary to perform meiotic recombination through double Holliday junctions. In yeasts (*Yarrowia lipolytica, Candida lusitaniae*, and *Pichia stipites*), a correlation has been observed between the importance of the GC trough near the centromeres (>10%) and the absence of MSH4/MSH5 genes^75^. It is therefore possible that the absence of this recombination pathway in *P. calceolata* induces more frequently double-strand break repair by synthesis-dependent strand annealing and a more rapid GC-biased conversion across the genome except at the centromeres where double-strand breaks are inhibited. This recombination inhibition may have important consequences on the evolution of the *P. calceolata* genome. Genes within low-GC regions are significantly longer and contain more introns than genes in other genomic regions. Because intron gain and loss are closely related to double-strand break repair and homologous recombination, we suggest that centromere genes retain more introns because double-strand breaks are reduced^83^. Variant analysis in *P. calceolata* populations could be targeted specifically in future studies to infer an estimation of recombination rate and more generally characterize the evolutionary processes controlling these large centromere regions.

### *P. calceolata* is one of the most abundant eukaryotes in the oceans

We have shown in this study that *P. calceolata* is cosmopolitan in oceanic samples above 10°C with a relative abundance generally >1% of all sequenced reads. In contrast to the coastal Pelagophyceae *A. anophagefferens* that can present high peaks of abundance^84^ no *P. calceolata* blooms are reported but *P. calceolata* is well-adapted to a large range of environmental conditions as suggested by previous studies^20,22^.

Although the abundance of an organism calculated from metabarcoding or metagenomic data provide only an indirect and relative quantification of organism abundances, both methods suggest that *P. calceolata* is one of the most abundant pico-nano eukaryote in offshore data. The high relative abundance of *P*.*calceolata* measured with a metabarcoding approach has recently been confirmed with a qPCR method^20^. In addition, we have shown that the metabarcoding approach probably underestimates the relative abundance of *P*.*calceolata* compared to the metagenomic analysis owing to the low copy number of rRNAs in this organism. Further studies may combine microscopic and flow sorting approaches with genomic data to assess the number of cells and the biomass of this organism in the oceans. Our modelling analysis has revealed a probable increase of *P. calceolata* relative abundance at the end of the century in high latitudes where the temperature is today too cold for this species. This result is coherent with previous studies suggesting a global increase of phytoplankton in subpolar regions^85,86^.

### Iron uptake and storage and modulation of iron needs

Iron is an essential element for the growth, photosynthesis, primary production, nitrogen fixation and reduction for PPEs^87^. Our results show that *P. calceolata* thrives in iron-poor waters and thus occupies a large ecological niche for a PPE. Two main strategies exist against iron deprivation: 1) optimisation of iron uptake and 2) modulation of iron needs. Optimisation of iron uptake does not seem to be the main strategy of *P. calceolata* in iron-poor environments. Genes coding for iron chelators and ferritin are absent from its genome, and gene coding for passive iron transporters are under- or equally represented compared to other PPEs. In contrast, phytotransferrins (ISIP2), putative iron storage proteins (ISIP3) and ferroportins are overrepresented in the *P. calceolata* genome. In Pelagophyceae, expression levels of ISIP genes are correlated with iron concentration at the global scale suggesting a transcriptomic regulation in response to iron availability^12^. Because phytotransferrins are dependent on carbonate ions, ocean acidification may reduce the efficiency of iron uptake in many species^88,89^. In consequence, organisms able to grow with very low iron concentration like *P. calceolata* may be favoured.

The presence of three ferroportin genes in *P. calceolata* is interesting since these transmembrane iron-export proteins play a major role in iron homeostasis in multicellular organisms^77^. Ferroportin function in micro-algae is unknown but could act to export iron from endosomes to the cytoplasm^90^. In the green alga *Chlamydomonas reinhardtii*, a ferroportin gene is overexpressed under low Fe conditions^91^. The function of the 3 ferroportin genes of *P. calceolata* could be investigated to understand their role in iron-poor environments. In addition, modulation of iron needs seems to be a major strategy for *P. calceolata*. All known molecular switches between ferrous and non-ferrous proteins are genetically possible in *P. calceolata* and the non-ferrous encoded protein genes FBA I, flaovodoxin and phytocyanin present a higher number of gene copies than the ferrous ones. It could be a sign that those switches are easier for *P. calceolata* than for other PPEs, but transcriptomic experiments are required to determine if this genetic overrepresentation has metabolic consequences.

### Nitrogen cycle

Expressing more than 90% of all nitrate transporter transcripts, Pelagophytes may dominate nitrate uptake and assimilation in the North Pacific Ocean^24^. Indeed, *P. calceolata* contains a large collection of genes for nitrate, nitrite and urea transporters. Interestingly, a few genes are ammonium transporters suggesting that the main source of nitrogen for *P. calceolata* is nitrite and/or nitrate. One remarkable feature of the *P. calceolata* genome is the presence of 3 genes carrying NIT domains (PF08376). This NIT domain was first described in bacterial nitrite and nitrate sensor proteins^92^. This sensor is an alpha-helical protein playing a signal transduction role regulating gene expression, cell motility and enzyme activity in *Klebsiella oxytoca*^93^. In pico-nano algae, NIT-sensing domains can be associated with a serine-threonine/tyrosine-kinase domain, suggesting signal transduction according to the concentration of intracellular nitrate/nitrite, or surrounded by 2 transmembrane domains suggesting extracellular sensing. Even though NIT-sensing domains can be found across various phyla, homologs of *P. calceolata* NIT proteins are restricted to Pelagophytes and Dictyochophytes. The NIT genes in *P. calceolata* are highly expressed in subtropical Pacific N-depleted waters, suggesting that these proteins have a role in transcription regulation according to nitrate availability^24^.

## Conclusion

Due to its widespread distribution and its high abundance in the open oceans, *Pelagomonas calceolata* can serve as an ecologically-relevant model to study marine photosynthetic protists. The chromosome-scale genome sequence, mostly telomere-to-telomere, generated in this study is an essential starting point for its detection in environmental datasets. This will allow studies of its ecological importance in the oceans at a molecular level. We have shown that the *P. calceolata* genome has specific genomic features potentially explaining its ecological success. The large repertoire of genes involved in nutrient acquisition from the environment is coherent with its widespread pattern of relative abundance distribution across different environments. The ecological niche of *P. calceolata* suggests that this alga will benefit from the global climate change with the extension of oligotrophic regions and global ocean warming. Future studies could use the *P. calceolata* genome to explore adaptation and acclimation processes controlling the distribution and abundance of this alga.

## Data availability

*Pelagomonas calceolata* genomic and transcriptomic reads, the genome assembly and gene prediction are available at the ENA (EMBL-EBI) website under the accession number PRJEB47931. Tara Oceans and Tara Polar Circle metagenomic sequences are archived at the ENA under the following accession numbers: PRJEB9740, PRJEB9691, PRJEB4352 and PRJEB1787.

## Acknowledgments

We thank the commitment of the following people who made this work possible: the Genoscope/CEA, the CNRS (in particular the Federation de Recherche R2022/Tara Oceans GO-SEE), Marie-José Garet-Delmas from the Roscoff Culture Collection for growing the RCC100 strain, Claude Scarpelli for support in high-performance computing. Computations were performed using the cobalt HPC machine. We acknowledge the financial support of FRANCE GENOMIQUE (ANR-10-INBS-09–08) and Oceanomics (ANR-11-BTBR-0008). We also thank the *Tara* Expedition Foundation and their partners for the organization of marine scientific expeditions (http://oceans.taraexpeditions.org). This article is contribution number XXX of *Tara* Oceans.

