## Supplementary Figures for "Genomic adaptation of the picoeukaryote *Pelagomonas calceolata* to iron-poor oceans revealed by a chromosome-scale genome sequence"

### Supplementary Figures of the manuscript: Genomic adaptation of the picoeukaryote *Pelagomonas calceolata* to temperate iron-poor oceans revealed by a chromosome-scale genome sequence.

Nina Guérin[1,2], Marta Ciccarella[1], Elisa Flamant[1,2], Paul Frémont[1,2], Sophie Mangenot[1,2], Benjamin Istace[1], Benjamin Noel[1], Sarah Romac[3], Charles Bachy[3], Martin Gachenot[4], Eric Pelletier[1,2], Adriana Alberti[1,2,5], Olivier Jaillon[1,2], Corinne Cruaud[1], Patrick Wincker[1,2], Jean-Marc Aury[1], Quentin Carradec[1,2]*

[1] Génomique Métabolique, Genoscope, Institut François Jacob, CEA, CNRS, Univ Evry, Université Paris-Saclay, 91057, Evry, France

[2]Research Federation for the Study of Global Ocean Systems Ecology and Evolution, R2022/Tara Oceans GO-SEE, 3 rue Michel-Ange, 75016, Paris, France

[3]Sorbonne Université, CNRS, Station Biologique de Roscoff, AD2M, UMR7144, Place Georges Tessier, 29680 Roscoff, France

[4]Sorbonne Université, CNRS, FR2424, Station Biologique de Roscoff, 29680 Roscoff, France

[5]Université Paris-Saclay, CEA, CNRS, Institute for Integrative Biology of the Cell (I2BC), 91198, Gif-sur-Yvette, France

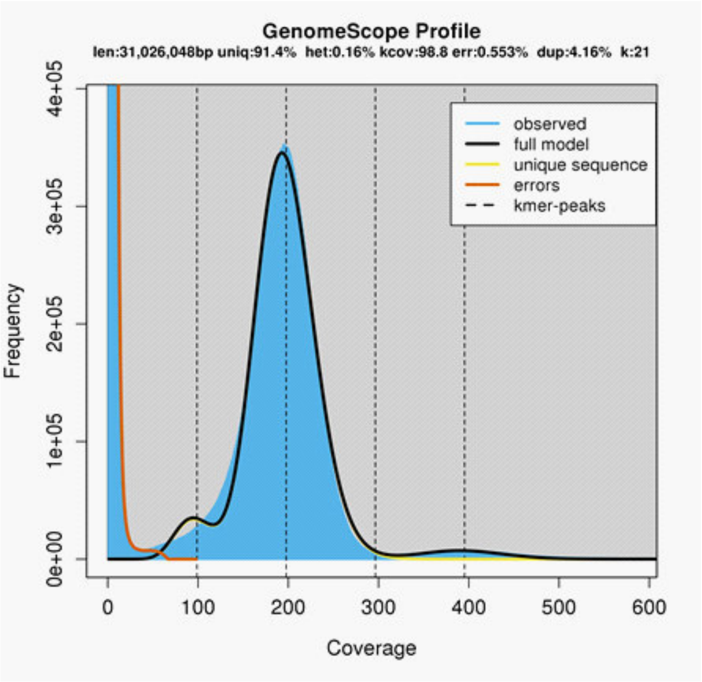


**Figure S1: K-mer profile of Illumina sequencing reads**.The graph shows the fit of the GenomeScope model (black) to the observed k-mer frequencies (blue). The k-mers presenting high frequency and low coverage (red) are considered as sequencing errors.

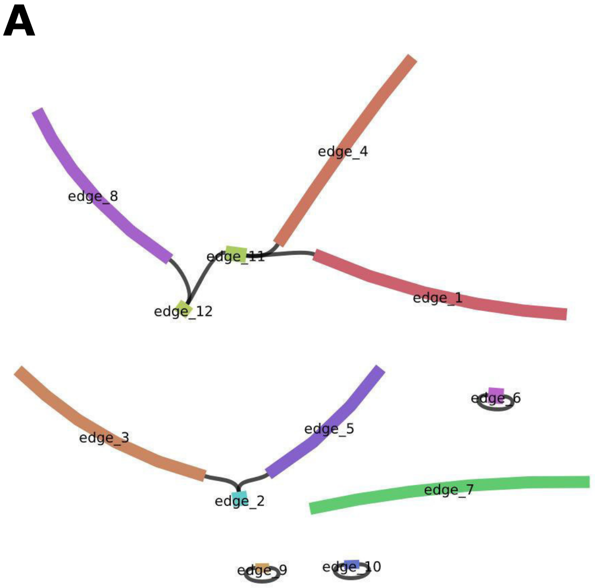


**B**

| Contig | Path | Size | Coverage |
|---|---|---|---|
| **1** | 11,1,* | 6 277 107 | 246 |
| **2** | *,7,* | 6 137 160 | 240 |
| **3** | *,-2,3,* | 5 383 939 | 241 |
| **4** | *,12,8,* | 5 311 685 | 243 |
| **5** | 11,4,* | 5 287 858 | 244 |
| **6** | *,5,2,* | 3 907 979 | 243 |
| **7** | 10 | 90 791 | 27 |
| **8** | 9 | 39 491 | 78 |

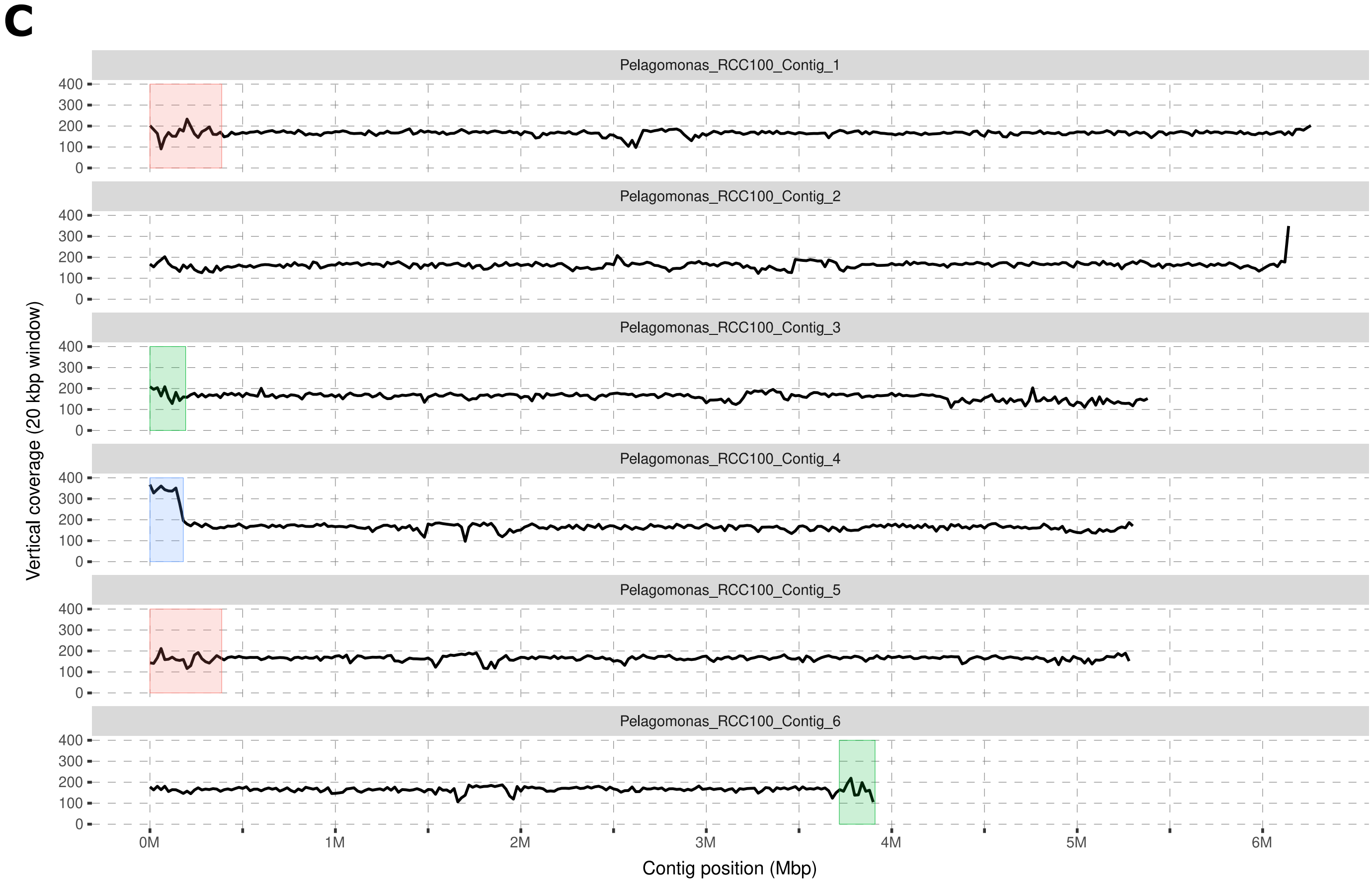


**Figure S2: Assembly and duplicated regions of the *P. calceolata* genome**. A) Flye assembly graph generated using bandage (Wick et al 2015). B) Table of the assembly paths and metrics of the assembly contigs detected by Flye. Numbers in the path column correspond to the edges in panel A. Telomeric repeats located at the ending edges of the assembly graph are reported as a star in the path column. C) Vertical coverage of Illumina reads in the *P. calceolata* genome. The dark line representes the number of Illumina short reads mapped over a window of 20 Kb on the 6 nuclear contigs of P. calcolata. Red and green boxes indicate highly similar regions (>99% of identity). The instability of the read coverage in these regions may be explained by the presence of variants in one of the two copies. The blue box is the only large region with a vertical coverage twice as high as the genome average.

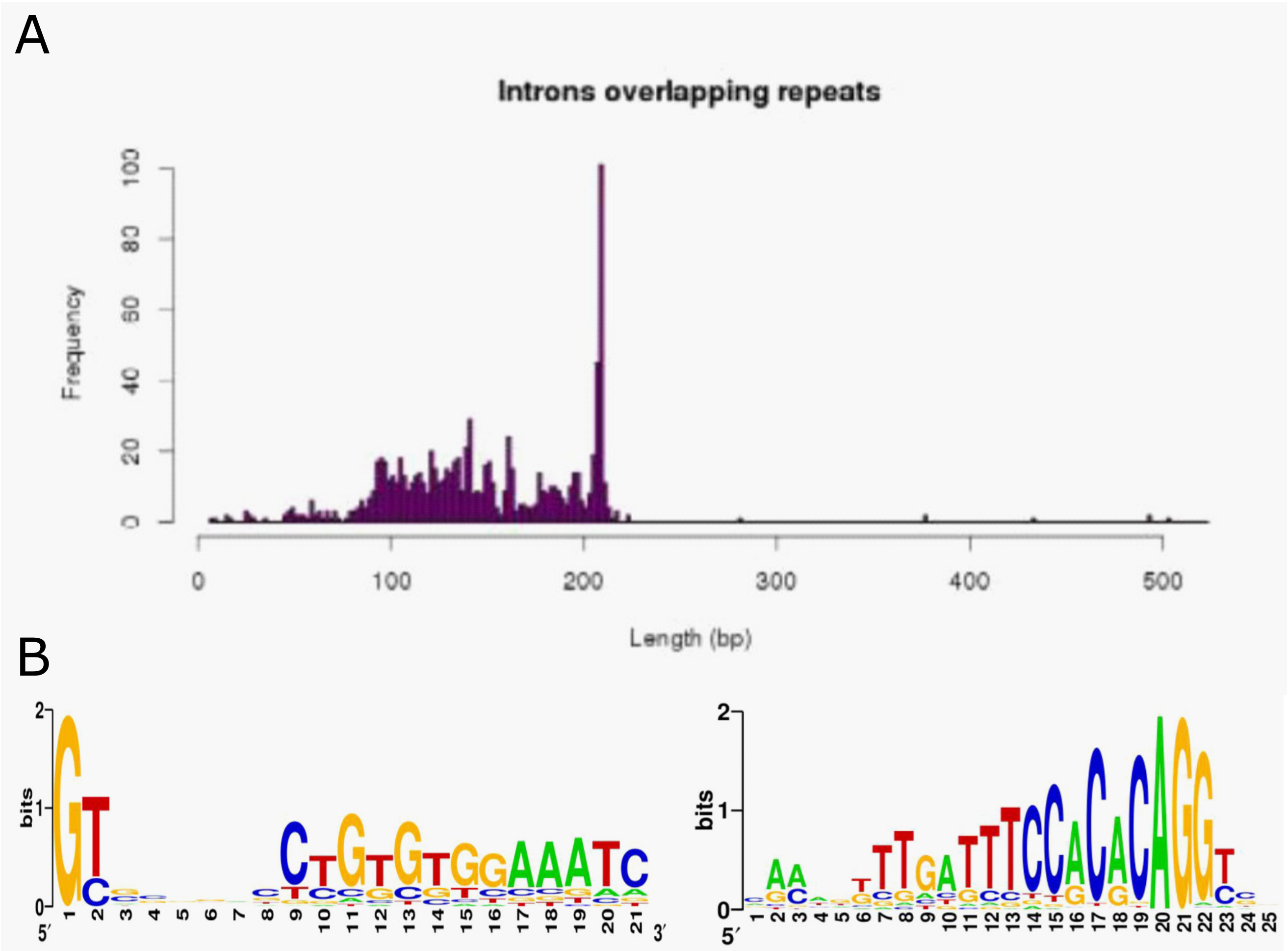


**Figure S3: Identification of Introner Elements from repeat families.** A) Distribution of putative IE lengths shows a pick around 200 bp. B) Logo representations of the starting and ending sequences of putative IE, revealing the presence of GT or GC donor splicing sites at the 5' ending, AG acceptor splicing site at the 3' ending, and conserved Terminal Inverted Repeats at the flanking regions.

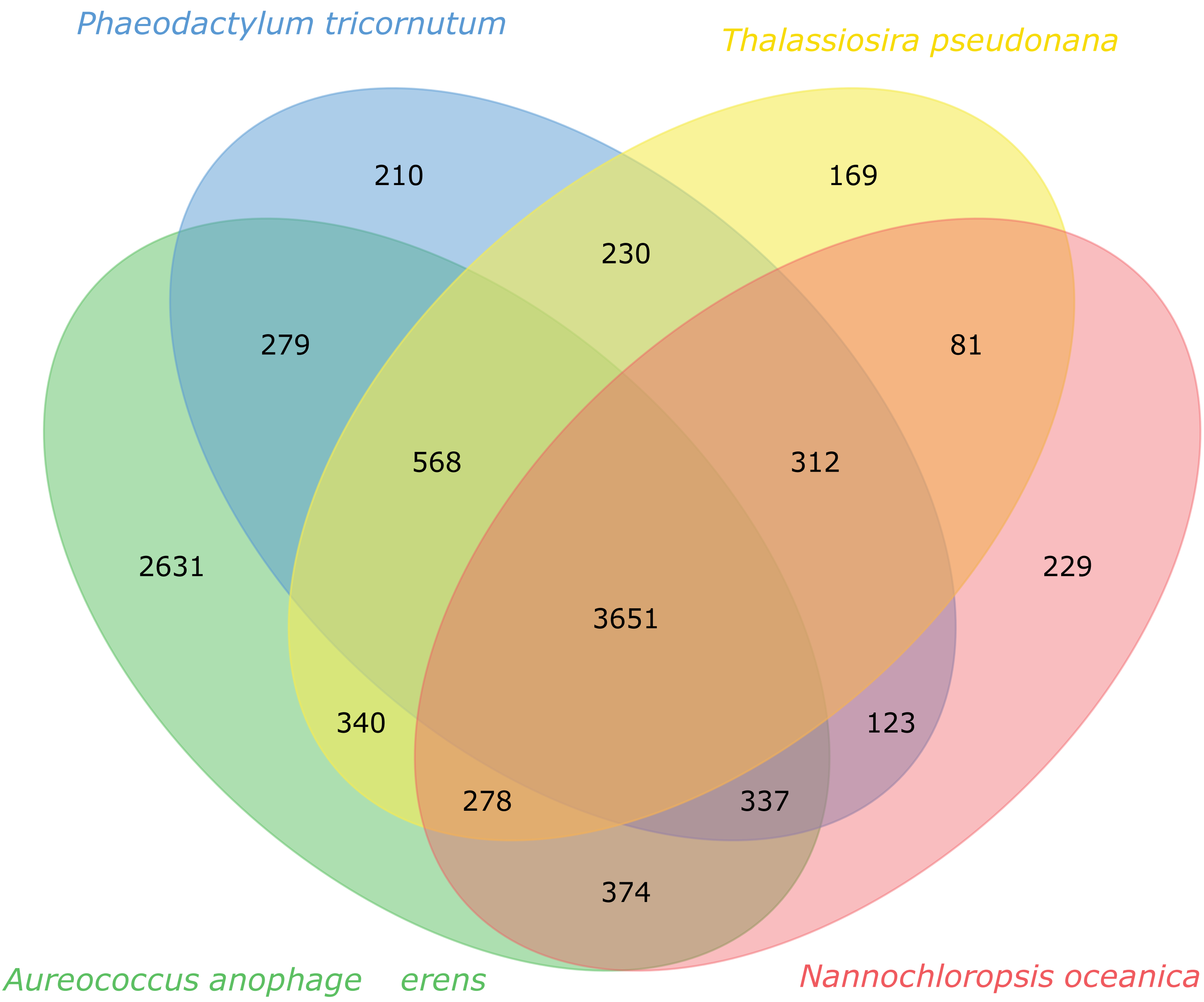


Figure S4. ***P. calceolata*** proteins shared with other Stramenopiles. The ve[illegible] displays the number of P. calceolata proteins homologous with at lea[illegible] another Stramenopile genome (alignment length >80% of the shortest pro[illegible] $<10^{-5}$)

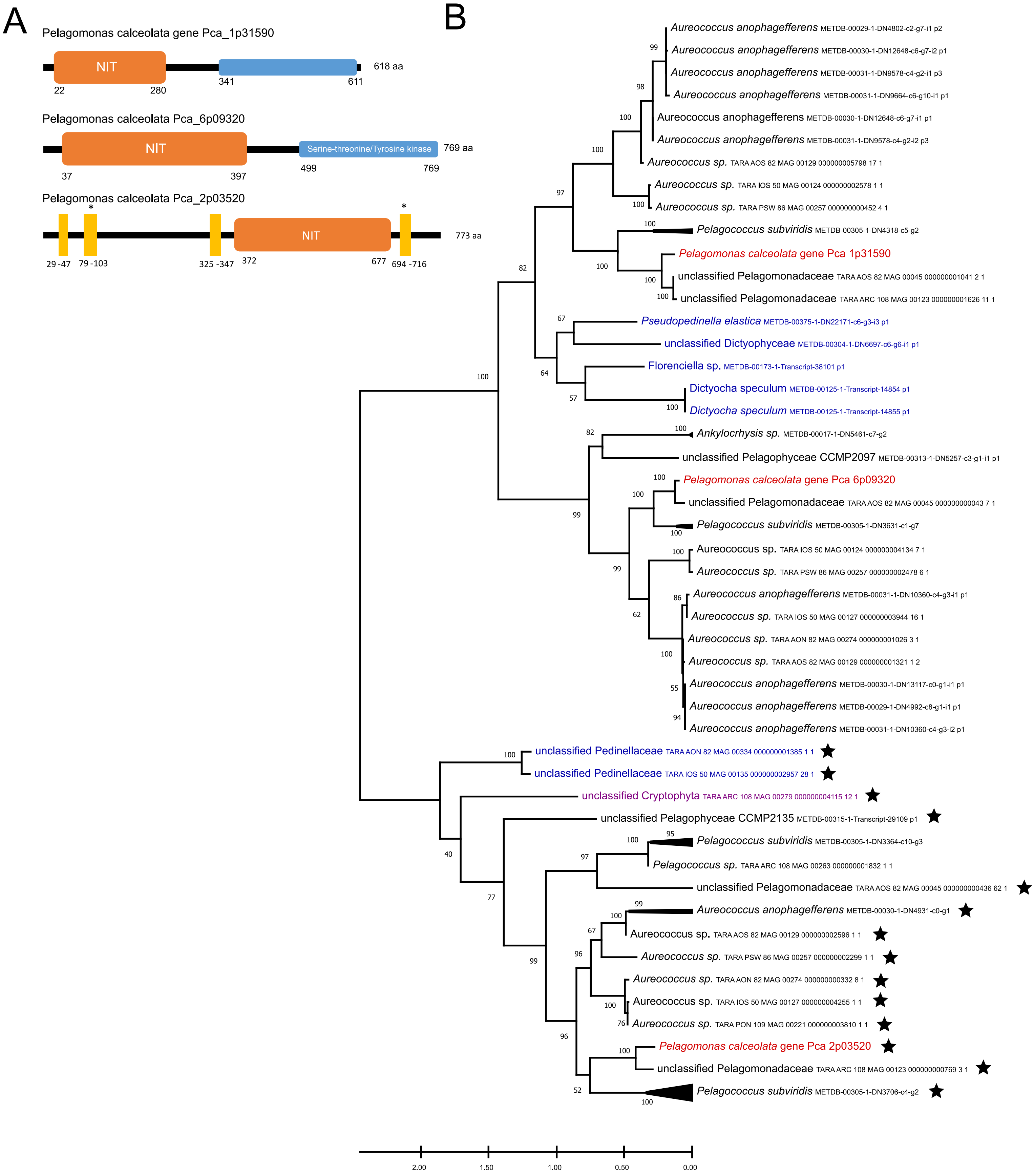


**Figure S5: Nitrate and nitrite sensing proteins similar in *P. calceolata* and other eukaryotes.** A) Domain organization of NIT-sensing proteins in *P. calceolata*. Orange boxes are NIT-sensing domains (IPR13587), blue boxes are serine-threonine/tyrosine kinase domain (IPR20635) and yellow rectangles are transmembrane domains. B) Maximum likelyhood phylogenetic tree made with JTT (Jones-Taylor-Thornton) model based on multiple alignment of NIT-sensing protein domains homologous to *P. calceolata*. Red: *P. calceolata* sequences. Black: other Pelagophyceae. Blue: Dictyophyceae. Purple: putative Cryptophyceae. The stars indicate NIT-sensing domain surrounded by transmembrane domains.

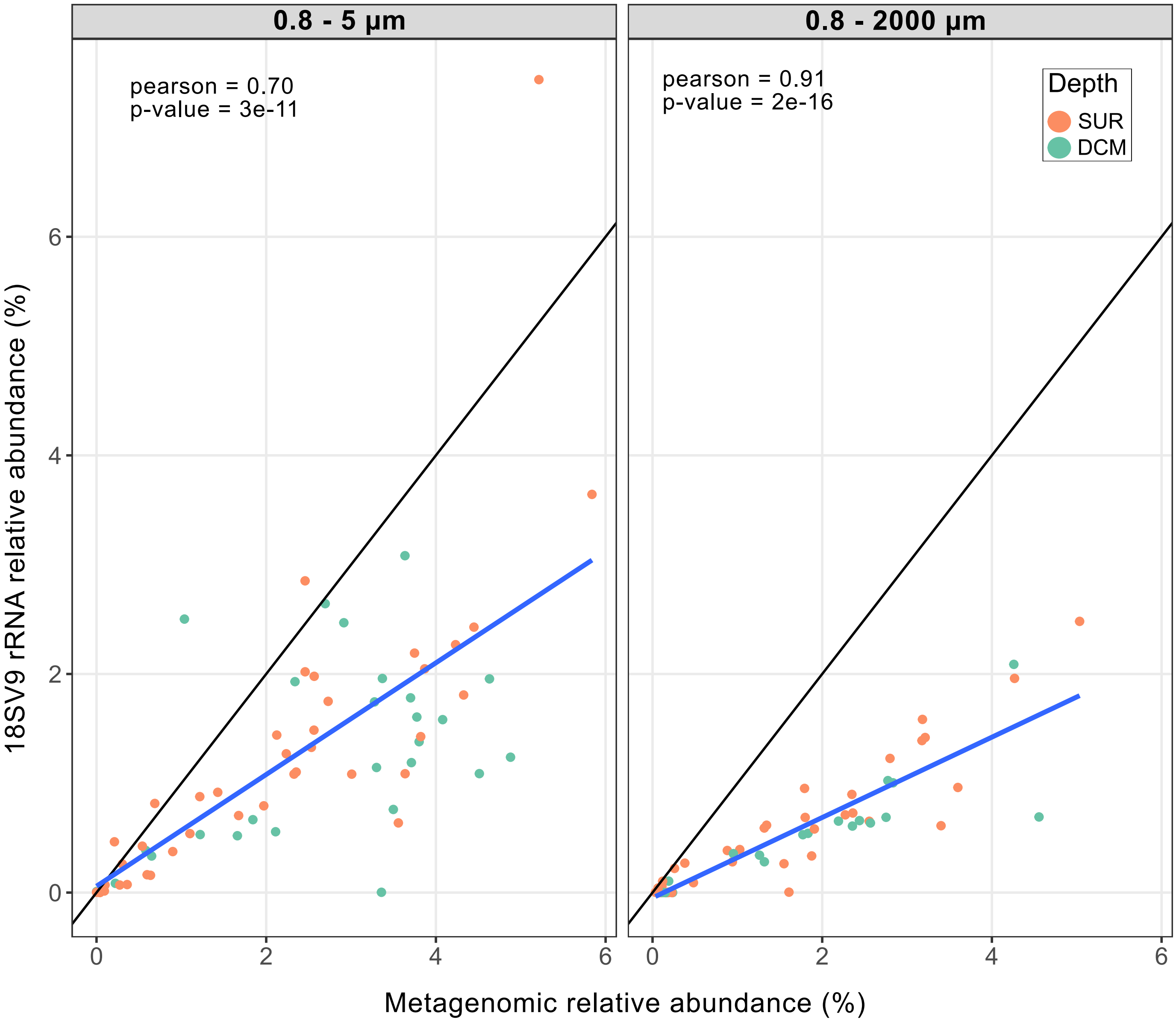


**Figure S6: Relative abundance of *P. calceolata* with 2 different methods.** Each dot represents a surface (orange) or DCM (green) sample. The relative abundance of *P.calceolata* is calculated from metagenomic reads aligned on the genome (x-axis) and with the amplified 18S rRNA sequence (y-axis) for size-fractions 0.8 - 5 µm (left) and 0.8 - 2000 µm (right). Blue lines are the linear regressions of observed values and black lines are expected trends.

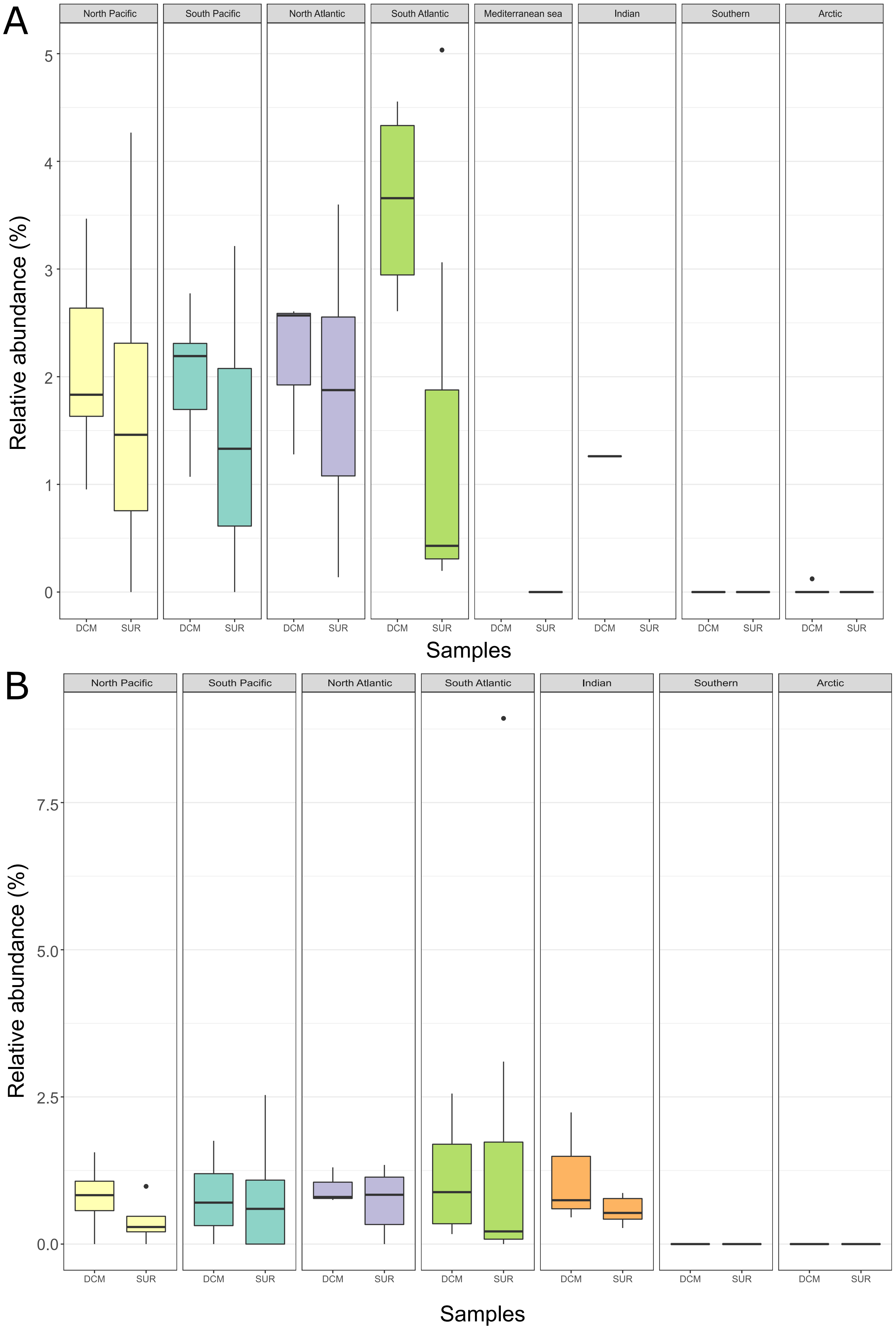


**Figure S7: Boxplot of the relative abundance of *P. calceolata* in each oceanic region in surface and DCM samples.** A) 0.2 - 3 µm size-fraction B) 0.8 - 2000 µm size-fraction.

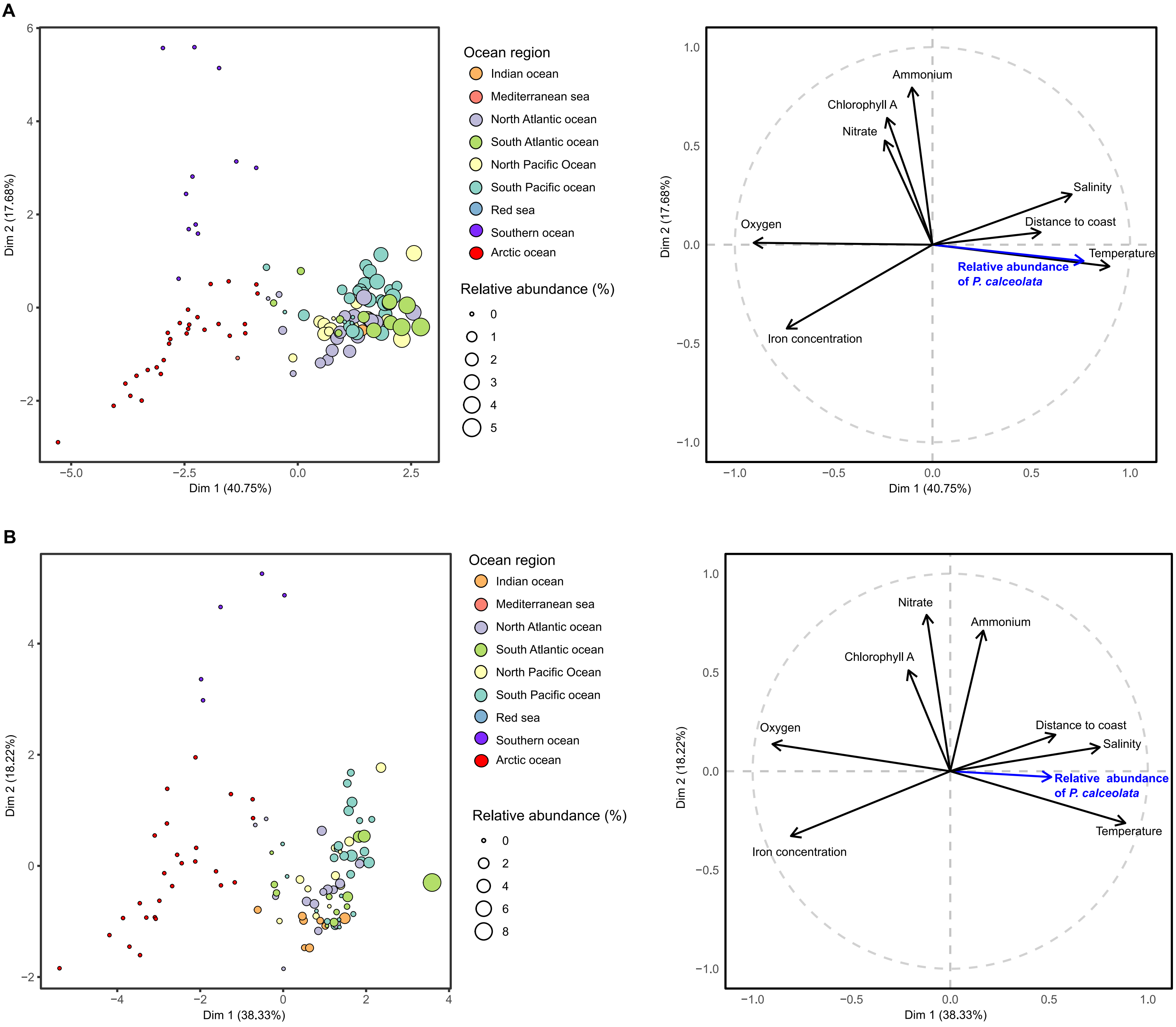


**Figure S8: Principal component analysis of the relative abundance of *P. calceolata* and the 8 environmental parameters.** A) 0.8-2000 µm size fraction and B) 0.2-3 µm size-fraction. Each dot represent an oceanic station with a size proportional to the relative abundance of *P. calceolata*. The colors indicate the oceanic basins.

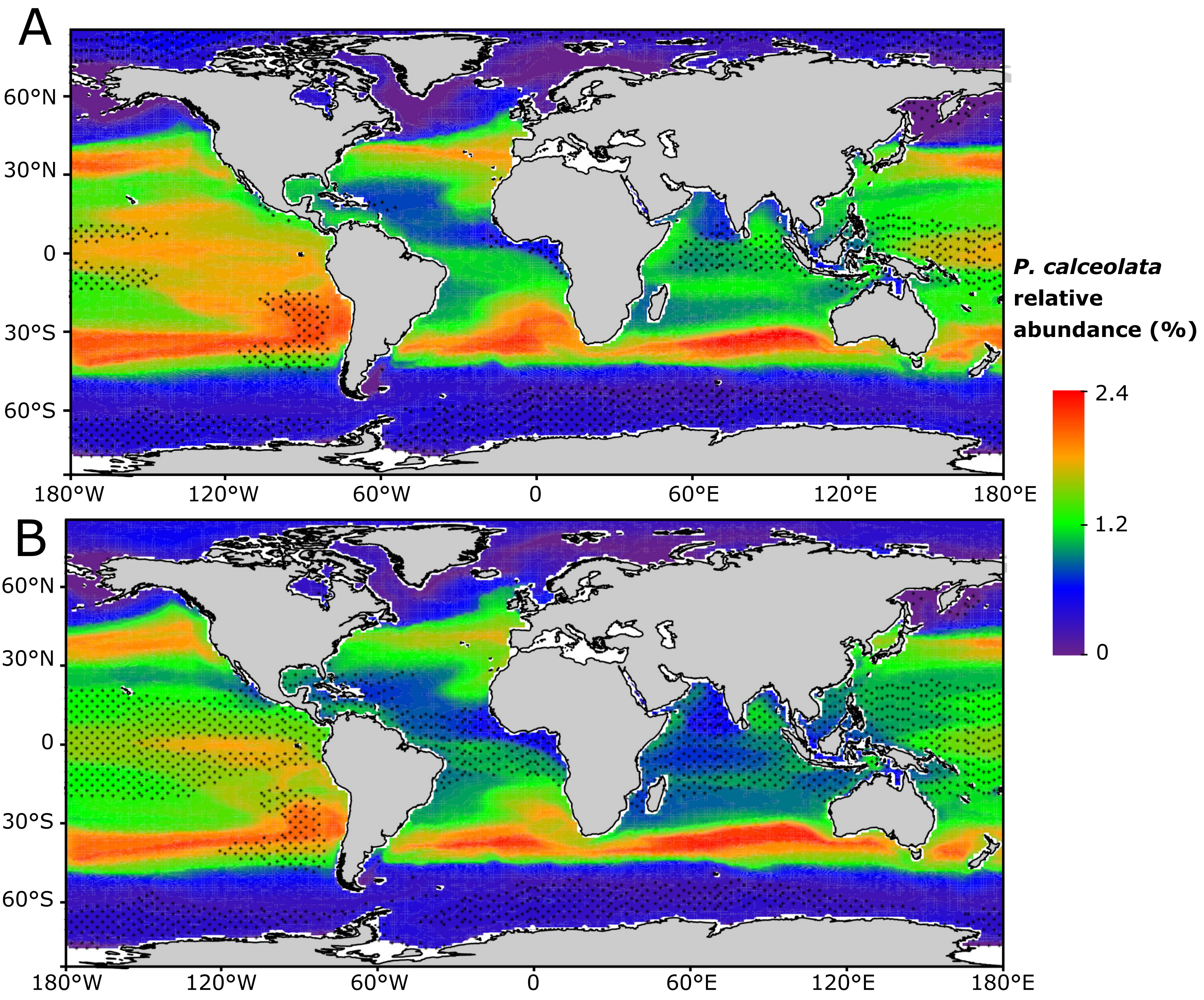


**Figure S9: Modelled genomic relative abundance of *P. calceolata* in present day (A) and at the end of the century (B).** The relative abundance of *P. calceolata* measured in Tara samples (0.8-2000 µm size-fraction) is modeled using the combination of 4 machine learning techniques trained on WOA18 environmental parameters at months, depths and sampling locations of Tara samples. Maps represent model projections on a multi-model mean of 6 Earth System Models in present day conditions (2006-15) (A) and end of the century conditions (2090-99) (B)
